# Influence of relay intercropping of barley with chickpea on biochemical characteristics and yield under water stress

**DOI:** 10.1101/2022.08.07.503101

**Authors:** Negin Mohavieh Assadi, Ehsan Bijanzadeh

## Abstract

Relay intercropping of legumes with cereals is a useful technique for yield improvement. Intercropping may be affected the photosynthetic pigments, enzymes activity and yield of barley and chickpea under water stress. To investigate the effect of relay intercropping of barley with chickpea on pigments content, enzymes activity and yield under water stress, a field experiment as split plot based on a randomized complete block design was conducted during 2017 and 2018. The treatments included irrigation regimes (normal irrigation and cutting off irrigation at milk development stage) as main plot. Also, cropping systems consisted of sole cropping of barley in December (b_1_) and January (b_2_), sole cropping of chickpea in December (c_1_) and January (c_2_), barley + chickpea in December (b_1_c_1_), barley in December + chickpea in January (b_1_c_2_), barley in January + chickpea in December (b_2_c_1_) and barley + chickpea in January (b_2_c_2_) as sub plot. Chlorophyll *a* content of barley increased in b_1_c_2,_ by consuming less water compared to sole cropping. In barley, sowing of barley in December intercropped with chickpea in December and January (b_1_c_1_ and b_1_c_2_) created a suitable canopy in pigment contents improvement. Late sowing of chickpea enhanced the carotenoid content of chickpea, catalase and peroxidase activities. Barley-chickpea intercropping reacted to water deficit through enzymes activity, water use efficiency and land equivalent enhancement compared to sole cropping. Under water stress, in b_1_c_2_, by increasing total chlorophyll and water use efficiency, grain yield of barley enhanced compared to b_1_. It seems that in b_1_c_2_, barley and chickpea response to water stress be increasing total chlorophyll and enzymes activity, respectively. In this relay intercropping treatment, each crop occupied and used the growth resources from different ecological niches at different times which is recommended in semi-arid areas.

## Introduction

Intercropping system is growing two or more plants in the same time and area [1,2,3]. In comparison to monoculture, intercropping mainly enhance the crop yield by more effective utilization of water resource and land [4, 5]. Relay intercropping describes a cropping system in which the life cycle of one crop overlaps that of another crop. Relay intercropping of legumes with cereals is a common practice in many regions due to its economic profitability, pest and weed suppression, high productivity and environmental protection [6]. Also, cereals-legumes relay intercropping system is widely practiced in areas where growing season is very short for two crops and rainfall is cut off during the active growth of the crop [7,8,9].

Barley (*Hordeum vulgare* L.) is the second main cereal and chickpea (*Cicer arietinum* L.) as third most important legume play an important role in agriculture of the world. After India, Turkey and Pakistan countries, Iran is the fourth biggest chickpea producer [10]. Barley intercropped with chickpea is one of the suitable kinds of intercropping systems in cool season of semi-arid areas for small farmers [11,12,13].

Water stress is one of the main limiting factors for crop production in semi-arid area via reducing water and nutrient uptake [14, 15]. The important challenge in intercropping is increasing the crop production by less water consumption [5]. Amanullah et al. [15] asserted that cereal-legume intercropping were the advantageous intercropping systems under water stress. However, crop species in intercropping may differ in their responses to growth under water shortage.

Photosynthetic pigments are sensitive to water stress extraordinarily which are main indicator of water deficit [16, 17]. Water stress accelerates the leaf senescence of crop and consequently chlorophyll content and photosynthesis rates reduced, negatively [18]. In *Arabidopsis thaliana*, chlorophyll content, chlorophyll fluorescence and RWC declined gradually under water stress [19]. It has been declared that under water stress, total chlorophyll content decreased because of degradation of chlorophyll *a* [17, 20]. To diminish the negative effect of water shortage on growth rate, plants have some defense mechanisms such as enhance antioxidant activities [21, 22]. Crops usually enhance the activity of catalase (CAT) and peroxidase (POD) in response to the water stress [23, 24]. Water stress triggers leaf water loss which declines relative water content (RWC) leads to growth rate reduction [25, 26].

In a common intercropping, the yield of each crop usually is less than that created in sole crop, while summation of relative yields often is higher than one [27, 28]. In contrast, some studies stated that intercropping create a significant yield advantage compared to monoculture [1, 29]. The yield of an intercropping system related to many factors such as sowing date, plant density, crop type, competition ratio and biotic and abiotic stress levels [30, 31, 32]. Barley-chickpea intercropping improved the sum yield in comparison to sole barley [13]. The interspecific interaction enhances the nutrient and water absorption from different depths of soil profile which caused yield enhancement [33]. Rahimi Azar et al. [34] suggested that the highest yield of chickpea was obtained from intercropping of chickpea with barley as 1:1 ratio. The chickpea seed yield was influenced by intercropping with barley, negatively [35].

One strategy to increase grain yield production of crop could be to change the sowing date and cropping system to withstand environmental stresses [36]. In relay intercropping, crop type and the sowing date of each component are crucial to final yield. Until now, no experiment has been carried out to consider the influence of relay intercropping of barley with chickpea on biochemical properties of each crop under water stress. It was hypothesized that relay intercropping influenced the photosynthetic pigment, antioxidant activities and yield of barley and chickpea under cutting off irrigation in the late season. The main goal of this field experiment was to detect the suitable relay intercropping system of barley-chickpea under water stress conditions and evaluate the biochemical changes of each crop in intercropping.

## Material and methods

### Filed experiment description and treatments

A consecutive two-year field experiment was laid out to evaluate the effect of late season water stress and different combinations of relay intercropping of chickpea with barley on some biochemical traits and yield. The experiment was conducted at College of Agriculture and Natural Resources of Darab (28°29’ N, 54°55’ E), Darab, Fars province, Iran during 2017 and 2018 growing seasons. The soil type in the experiment site was loam (fine, loamy, carbonatic, hyperthermic, typic Torriorthents) and the other soil characteristics are given in Table 1. The study area has a semi-arid climate with cool and rainy winters and dry and hot summers. Likewise, weather data for the research field during the both years are presented in Table 2.

**Table 1.** Some physical and chemical characteristics of the soil in the experimental site (depth of 0-30 cm) during 2017 and 2018 growing seasons.

| Year | Sand (%) | Silt (%) | Clay (%) | Organic content (%) | Electrical conductivity (dS m<sup>-1</sup>) | pH | N (%) | P (mg kg<sup>-1</sup>) | K (mg kg<sup>-1</sup>) |
| --- | --- | --- | --- | --- | --- | --- | --- | --- | --- |
| 2017 | 38.01 | 43.95 | 18.04 | 0.821 | 1.011 | 7.31 | 0.075 | 51 | 311 |
| 2018 | 38.12 | 44.21 | 17.67 | 0.843 | 1.015 | 7.33 | 0.074 | 52 | 307 |

**Table 2.** Minimum and maximum air temperatures, monthly rainfall, and pan evaporation of the experimental site during 2017 and 2018 growing seasons.

| Month | Temperature (°C) |  |  |  |  |  | Rainfall<br>(mm) |  | Pan evaporation<br>(mm) |  |
| --- | --- | --- | --- | --- | --- | --- | --- | --- | --- | --- |
|  | 2017-18 |  |  | 2018-19 |  |  | 2017-18 | 2018-19 | 2017-18 | 2018-19 |
|  | Min | Max | Mean | Min | Max | Mean |  |  |  |  |
| December | 5.1 | 20.1 | 12.6 | 5.7 | 21.2 | 13.5 | 2.2 | 24.8 | 77.6 | 88.4 |
| January | 4.4 | 19.2 | 11.8 | 3.8 | 23.4 | 13.6 | 94.2 | 78.3 | 66.3 | 78.5 |
| February | 8.6 | 21.4 | 15.0 | 9.2 | 23.6 | 17.9 | 44.3 | 38.9 | 88.1 | 91.2 |
| March | 10.5 | 25.4 | 18.0 | 8.1 | 28.5 | 19.3 | 99.5 | 22.1 | 143.1 | 175.3 |
| April | 12.9 | 29.8 | 21.4 | 8.9 | 33.2 | 22.6 | 0.0 | 0.0 | 202.3 | 218.9 |
| May | 17.9 | 34.4 | 26.2 | 15.8 | 38.1 | 30.0 | 0.0 | 0.0 | 276.2 | 297.4 |
| Total |  |  |  |  |  |  | 240.2 | 164.1 | 853.6 | 949.7 |

The experiment in each year was conducted as split plot based on a randomized complete block design with three replications. The treatments included irrigation regimes in two levels (normal irrigation and cutting off irrigation at full anthesis stage of barley [Zadoks growth stage (ZGS69)] [37] were as main plot. Cropping system treatments as sub plot consisted of sole cropping of barley in December (b_1_) and January (b_2_), sole cropping of chickpea cultivar in December (c_1_) and January (c_2_), and different combinations of intercropping consisted of intercropping of barley + chickpea in December (b_1_c_1_), intercropping of barley in December + chickpea in January (b_1_c_2_), intercropping of barley in January + chickpea in December (b_2_c_1_) and intercropping of barley + chickpea in January (b_2_c_2_). The seeds of barley (Zehak cultivar) and chickpea (Darab cultivar) were provided from the Agriculture and Natural Resources Research Center of Darab, Fars Province, Iran. Zehak is a six-rowed barley cultivar with a medium plant height 50-80 cm. The growing season length for Zehak is approximately 150 days, which is specifically adapted to growing in the warm and arid regions of Iran [38]. Also, Darab is an early mature cultivar of chickpea with a plant height of 28 cm, have a semi-upright position, which is suitable for hot and dry areas [39].

The plot size was 3m×2m where was surrounded with a 30 cm high earth berm, by a 1m wide buffer space between the plots. Seedbed was prepared by mouldboard ploughing and disking. Then, uniform seeds were sown handily at a soil depth of 3 cm in rows 30 cm apart, giving 250 and 40 plants m^-2^ planting density for barley and chickpea, respectively. The barley and chickpea were designed as replacement series with a ratio of one row barley and one row chickpea in intercropping plots. Based on soil test (Table 1), phosphorus (P) as superphosphate triple source at the rate of 50 kg ha^-1^ and Nitrogen (N) as urea source at the rate of 60 kg ha^-1^ were used in the field experiment. Total P and half dose of N used in the soil at sowing and remaining half N dose used by irrigation water at beginning of the stem elongation stage of barley (ZGS31). The seeds were sown on December 15^th^ or January 15^th^, based on sowing dates of cropping systems.

The soil water content was traced in each plot at 30 cm depth down to 90 cm, gravimetrically. Before each irrigation, the soil profile was sampled up to 90-cm by an auger. Then, the volume of water applied in normal irrigation was accorded to restoring root zone moisture deficit (when 50% of available water was depleted in effective root-zone depth of 90 cm) to near-field capacity [26]. A surface drip irrigation system was applied for irrigation. A 20 mm diameter polyethylene pipe with in-line drippers at 40 cm intervals was placed on one side of each planting row. Five irrigations were applied during the crop season for normal irrigation and three irrigations were applied for cutting off irrigation regime at full anthesis stage (ZGS69) of barley. Total water applied (m^3^) (irrigation amount + rainfall) in each irrigation regime and cropping system in 2017 and 2018 growing seasons are presented in Fig. 1. To determine the photosynthetic pigments, antioxidant enzyme activity and RWC the top leaf of barley and chickpea were sampled at end of milk development stage of barley (ZGS 77). At the crop maturity stage on May, the plants in the central 1 m^2^ of each plot were hand harvested manually. Then, the samples were oven dried at 72 °C for 48 h and weighted for biological yield. Finally, the samples were threshed and grains were separated and weighed for grain yield determination.

**Figure 1.**
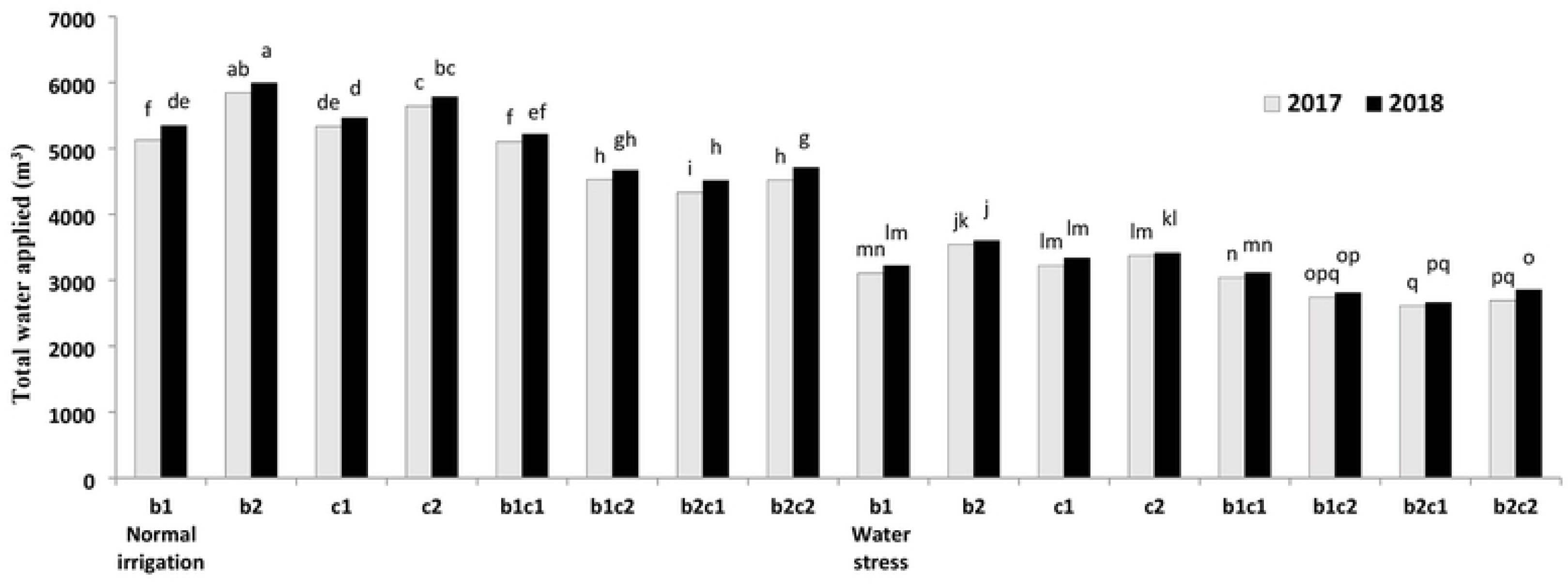
Total water applied (m3) in each irrigation regime and cropping system during the *two* growing seasons. b,: Sole cropping of barley in December b_2_: Sole cropping of barley in January, c_1_: Sole cropping of chickpea in December c_1_: Sole cropping of chickpea in January, b_1_c_1_: lntcrcropping of barley+ chickpea in December; b,c,, lntercropping of barley in December+ chickpea in January, b_2_c_1_: lntercropping of barley in January+ chickpea in December, b_1_c_1_: lntercropping of barley + chickpea in January. Means sharing the same letter do not differ significantly based on LSD (pSO.OS) test.

### Chlorophyll and carotenoid content assessment

The chlorophyll content was measured by fresh tissue of top leaf in each plot. Ten ml of 80% acetone was added to 200 mg of leaf tissue gradually and ground by a mortar and pestle. The created slurry was centrifuged for 10 min at 4000 rpm, and the supernatant was filtered through Whatman No. 2 filter paper placed in a funnel as the solution was transferred. Absorbance was measured by a double-beam UV-VIS spectrophotometer (UV-1900 spectrophotometer, Shimadzu, Japan) at λ = 645, 663, and 470 nm. Chlorophyll *a*, *b* and total and Carotenoid was calculated according to the following equations [40]:

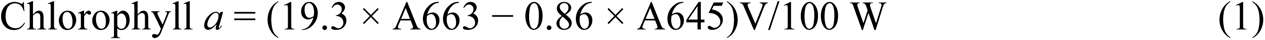

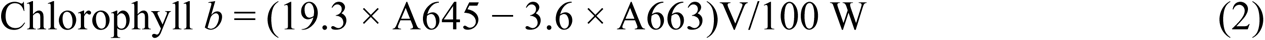

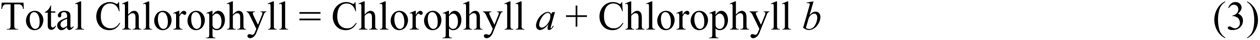

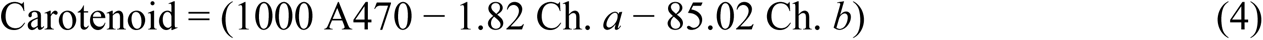

Where, V is volume of purified solution, W is leaf fresh weight, and A663, A645, and A470 were optical absorption wavelengths at 663, 645, and 470 nm, respectively.

### Antioxidant enzymes assay

For enzyme extraction, 0.5 g of fresh leaves were ground to fine powder in liquid nitrogen by mortar and pestle and then homogenized in 2 mL extraction buffer containing 10% (w/v) polyvinyl pyrrolidone (PVP) in 50 mM potassium-phosphate buffer (pH 8), 1 mM dithiothreitol (DTT), and 0.1 mM ethylene diamine tetra acetic acid (EDTA). The homogenate was centrifuged at 20,000×g (4°C) for 30 minutes. The supernatant was used to assess antioxidant enzymes of catalase and peroxidase.

### Catalase enzyme

The catalase enzyme activity (CAT) was determined using spectrophotometer (UV-160A) according to the method of Aebi [41], with monitoring the decrease in absorbance at 240 nm because of H_2_O_2_ consumption. One mL of reaction mixture contained 50 mM potassium phosphate buffer (pH= 7.0) and 15 mM H_2_O_2_. The reaction was initiated with adding 50μL of crude extract to this solution. CAT activity was expressed as units (μmol H_2_O_2_ consumed per minute) per milligram of protein.

### Peroxidase enzyme

The peroxidase enzyme activity (POD) was evaluated by method of Chance and Maehly [42]. One mL of reaction mixture contained 13 mM guaiacol, 50 mM potassium phosphate buffer (pH 7) and 5 mM H_2_O_2_. Increase in absorbance because of oxidation of guaiacol (extinction coefficient: 26.6 mM^- 1^.cm^- 1^) was traced at 470 nm for a minute. Peroxidase activity was expressed as units (μmol guaiacol oxidized per minute) per milligram of protein.

### Leaf relative water content

The leaf relative water content (RWC) was measured by method of Machado and Paulsen [43]. Eight leaf discs (8 mm in diameter) from fully expanded of flag leaf were weighed for determination of fresh weight (FW). The leaf discs were kept in distilled water for 6 h, then dried with filter paper and weighed for determination of total weight (TW). Dry weight (DW) was determined after drying the discs at 70 °C in oven for 24 h. Finally, the RWC was determined as:

### RWC= [(FW-DW)/(TW-DW)] ×100

### Water use efficiency

The water use efficiency (WUE) in each treatment was evaluated as the ratio of seed yield (g. m^- 2^) to total water consumed (mm) [44].

### Land equivalent ratio

The land equivalent ratio (LER) is expressed as the land equivalent required for growing either crop in intercropping compared to the land area required to sole cropping of each crop. The LER total (LER_t_) were calculated as [45]:

LERt= LERb+ LERc

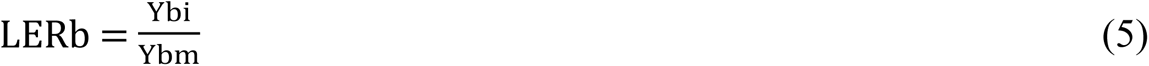

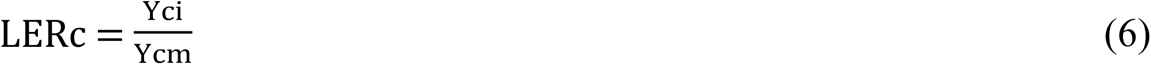

Where LER_b_ and LER_c_ were land equivalent ratios of barley and chickpea, respectively; Y_bi_ m and Y_ci_ were the yields of barley and chickpea in monoculture; and Ybi and Yci were the yields of barley and chickpea in intercropping system. When the LER_t_ value was more than one, intercropping was more useful as compared to sole cropping. Controversy, when the LER was less than one, intercropping affected yield of crops, negatively [6].

### Competition ratio

The competition ratio (CR) is a suitable index to determine the competitive ability between two crops in intercropping. CR shows stronger competitive ability to the species and is more beneficial compared to other indices. The CR was determined by the following equations [6] :

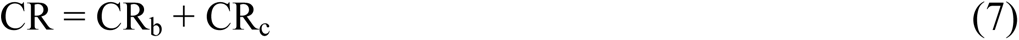

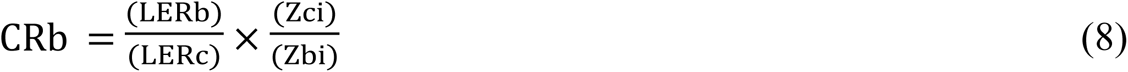

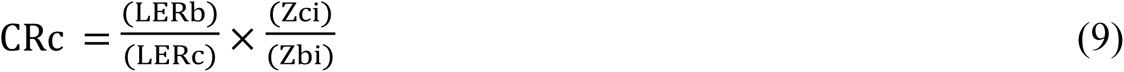

Where Z_bi_ is the sown proportion of barley intercropped with chickpea and Z_ci_ is the sown proportion of chickpea intercropped with barley. CR_b_ is the competition ratio of barley and CR_c_ is the competition ratio of chickpea.

### Statistical analyses

Data was analyzed by SAS software 2012 (version 9.4) and the means were compared using the least significant differences (LSD) test at 0.05 probability level (p≤ 0.05). Because of significant effect of year × irrigation regime × cropping system on considered traits, the data of two years for barley and chickpea presented.

## Results

### Analysis of variance

Results of combined analysis of variance over years demonstrated that the main effect of year was significant on all of the considered traits at 0.05 probability level (Table 3). It might be related to different temperature, rainfall and evaporation during the active growth of barley and chickpea (Table 2). Also, the interaction effects of year × irrigation regime × cropping system was significant (p ≤ 0.05) on total water applied, chlorophyll a, chlorophyll b, carotenoid, catalase and peroxidase of barley and chickpea (Table 3). On the other hand, this interaction had a significant effect at %5 probability level on relative water content, yield attributes, water use efficiency and competition indices (Table 4).

**Table 3.** Combined analysis of variance for total water applied, pigment content and enzyme activity of barley and chickpea.

| Source of variance (S.O.V) | Degrees of freedom (df) | Mean squares (MS) |  |  |  |  |  |  |  |  |  |  |  |  |  |
| --- | --- | --- | --- | --- | --- | --- | --- | --- | --- | --- | --- | --- | --- | --- | --- |
|  |  | TWA |  | Chl. *a* |  | Chl. *b* |  | CAR |  | Total Chl. |  | CAT |  | POX |  |
|  |  | Barley | Chickpea | Barley | Chickpea | Barley | Chickpea | Barley | Chickpea | Barley | Chickpea | Barley | Chickpea | Barley | Chickpea |
| Year(Y) | 1 | 12.5** | 10.33* | 0.76* | 1.21* | 0.13* | 1.56* | 3.96 * | 14.56* | 5.32* | 11.96* | 5.86* | 4.63* | 5.36* | 2.36* |
| Replication (R) | 4 | 0.025<sup>ns</sup> | 0.066<sup>ns</sup> | 0.0088<sup>ns</sup> | 0.086 <sup>ns</sup> | 0.0002 <sup>ns</sup> | 0.0063<sup>ns</sup> | 0.0031 <sup>ns</sup> | 0.036 <sup>ns</sup> | 0.0061 <sup>ns</sup> | 0.065 <sup>ns</sup> | 0.0073 <sup>ns</sup> | 0.0036 <sup>ns</sup> | 0.086 <sup>ns</sup> | 0.031 <sup>ns</sup> |
| Irrigation regime (I) | 1 | 7.65* | 11.26* | 0.027* | 0.056* | 0.0178* | 0.189* | 0.014<sup>ns</sup> | 0.086 <sup>ns</sup> | 0.167* | 0.986* | 0.351* | 0.236* | 0.498* | 0.569<sup>ns</sup> |
| Cropping system (C) | 5 | 4.36* | 3.68* | 0.0921** | 0.653* | 0.0309** | 0.410* | 0.0013<sup>ns</sup> | 0.013* | 0.052* | 0.056* | 0.098<sup>ns</sup> | 0.088* | 0.103* | 0.201<sup>ns</sup> |
| I×C | 5 | 11.57* | 14.63 <sup>ns</sup> | 0.00768 <sup>ns</sup> | 0.00563* | 0.0023* | 0.036 <sup>ns</sup> | 0.0031* | 0.036 <sup>ns</sup> | 0.0136* | 0.0189* | 0.0436 <sup>ns</sup> | 0.0365 <sup>ns</sup> | 0.891 <sup>ns</sup> | 0.0986* |
| Y×I | 1 | 23.45 | 12.98* | 0.00019 <sup>ns</sup> | 0.00086 | 0.000046 <sup>ns</sup> | 0.0021 | 0.00018<sup>ns</sup> | 0.0025 | 0.00065 <sup>ns</sup> | 0.0658 | 0.00087 | 0.0035 | 0.0023 | 0.0064 |
| Y×C | 5 | 2657.3 | 5684.3* | 0.000023<sup>ns</sup> | 0.000043 | 0.000021 <sup>ns</sup> | 0.00613 | 0.00015 <sup>ns</sup> | 0.0014 | 0.00046 <sup>ns</sup> | 0.045 | 0.000086 | 0.000011 | 0.00095 | 0.00068 |
| Y×I×C | 5 | 456879.3* | 436357.3* | 0.00061* | 0.00096* | 0.0003* | 0.0032* | 0.00031* | 0.00894* | 0.00025* | 0.00035* | 0.00032* | 0.00057* | 0.0092* | 0.0035* |
| Error | 35 | 10.33 | 9.66 | 0.002433 | 0.002369 | 0.002136 | 0.03561 | 0.00463 | 0.0261 | 0.0007862 | 0.008661 | 0.009823 | 0.006357 | 0.01235 | 0.01276 |
| CV% |  | 4.87 | 11.23 | 4.68 | 5.39 | 5.23 | 4.98 | 3.36 | 11.69 | 6.11 | 12.37 | 5.11 | 3.98 | 6.54 | 7.93 |

TWA: total water applied; Chl. *a*: Chlorophyll *a*; Chl. *b*: Chlorophyll *b*; CAR: Carotenoid; Total chl.: Total chlorophyll; CAT: Catalase; POX: Peroxidase; LER: land equivalent ratio; CR: Competition ratio. <sup>ns</sup>, \* and \*\*: no significant and significant at the 5% and 1% probability levels, respectively.

**Table 4.** Combined analysis of variance for relative water content, yield attributes, water use efficiency and competition indices of barley and chickpea.

| Source of variance (S.O.V) | Degrees of freedom (df) | Mean squares (MS) |  |  |  |  |  |  |  |  |  |  |  |  |  |
| --- | --- | --- | --- | --- | --- | --- | --- | --- | --- | --- | --- | --- | --- | --- | --- |
|  |  | RWC |  | Gy |  | By |  | HI |  | WUE |  | LER |  | CR |  |
|  |  | Barley | Chickpea | Barley | Chickpea | Barley | Chickpea | Barley | Chickpea | Barley | Chickpea | Barley | Chickpea | Barley | Chickpea |
| Year(Y) | 1 | 23.44 * | 12.36* | 5727.20* | 3266.3* | 6312.32* | 12386.51* | 11.31* | 10.91* | 28.93* | 98.32* | 14.33* | 9.53* | 4.32* | 2.36* |
| Replication (R) | 4 | 13.91<sup>ns</sup> | 15.69<sup>ns</sup> | 96.32<sup>ns</sup> | 54.23<sup>ns</sup> | 862.39<sup>ns</sup> | 563.28<sup>ns</sup> | 9.88<sup>ns</sup> | 15.36<sup>ns</sup> | 15.68<sup>ns</sup> | 10.41<sup>ns</sup> | 12.44<sup>ns</sup> | 2.64<sup>ns</sup> | 1.31<sup>ns</sup> | 4.54<sup>ns</sup> |
| Irrigation regime (I) | 1 | 655.32** | 866.34* | 6821.37* | 7433.7* | 2569.96** | 1256.38* | 154.32* | 186.76* | 986.33 | 864.38** | 5.63* | 4.36* | 2.96* | 1.76* |
| Cropping system (C) | 5 | 684.23* | 684.23* | 39154.51* | 22368.5* | 7568.64* | 6354.28* | 863.94<sup>ns</sup> | 602.31* | 765.35* | 625.97<sup>ns</sup> | 15.3* | 10.91* | 9.36* | 10.37<sup>ns</sup> |
| I×C | 5 | 244.36* | 278.39<sup>ns</sup> | 557.11* | 453.68* | 235.04* | 133.98<sup>ns</sup> | 431.22* | 45.362<sup>ns</sup> | 153.68<sup>ns</sup> | 25.63<sup>ns</sup> | 15.31<sup>ns</sup> | 4.36<sup>ns</sup> | 2.31<sup>ns</sup> | 3.33* |
| Y×I | 1 | 11.86<sup>ns</sup> | 50.36* | 2523.4<sup>ns</sup> | 2438.71* | 3111.37<sup>ns</sup> | 7423.69* | 17.44* | 10.76* | 52.93<sup>ns</sup> | 45.36* | 11.91<sup>ns</sup> | 12.32* | 10.69* | 9.32* |
| Y×C | 5 | 27.36<sup>ns</sup> | 38.91* | 1589.3<sup>ns</sup> | 1025.96<sup>ns</sup> | 1457.56<sup>ns</sup> | 1598.78* | 11.59* | 59.67<sup>ns</sup> | 11.21<sup>ns</sup> | 10.86* | 11.56* | 10.91<sup>ns</sup> | 7.43<sup>ns</sup> | 6.54<sup>ns</sup> |
| Y×I×C | 5 | 5.33* | 7.36* | 22403.2* | 28456.81* | 4222.44* | 4203.56* | 8.80* | 45.36** | 43.96* | 57.48* | 22.33* | 9.86* | 5.63* | 4.36* |
| Error | 35 | 1.63 | 0.56 | 277.3 | 156.98 | 399.48 | 766.2 | 212.3 | 127.86 | 14.68 | 10.81 | 1.23 | 2.38 | 1.54 | 2.38 |
| CV% |  | 7.86 | 9.35 | 10.58 | 12.36 | 11.21 | 10.43 | 12.34 | 10.41 | 8.51 | 12.29 | 4.23 | 5.11 | 6.11 | 7.57 |

RWC: relative water content; Gy: grain yield, By: biological yield; HI; harvest index; WUE: water use efficiency; LER: land equivalent ratio, CR: competition ratio. ns, \* and \*\*: no significant and significant at the 5% and 1% probability levels, respectively.

### Weather conditions and total water applied

Throughout the growth period of barley and chickpea from December to May, the mean temperature in 2018 growing season was more that 2017 (Table 2). In each year, the maximum temperatures were registered in April-May and the minimum in December- January. By increasing the mean temperature, the water evaporation in second year (949.7 mm) increased compared the first year (853.6 mm). Also, total rainfall during the active growth stages of 2017 and 2018 was 240.2 and 164.1 mm, respectively. A significant portion of rainfall occurred in January (94.2 mm) and March (99.5 mm) of the first year (Table 2). During the 2017 and 2018, in two irrigation regimes, intercropping treatments consumed less total water (irrigarion amount+rainfall) compared to sole cropping. On the other hand, in two years, there was no efficient rainfall in April to May when the water demand of barley and chickpea enhanced sharply to complete the grain filling period (Table 2). A significant difference (p≤ 0.05) was obtained between total water applied of relay intercropping of barley with chickpea (b_1_c_2_, b_2_c_1)_ and simultaneous intercropping in January (b_2_c_2_) with sole cropping treatments (Fig. 1). In 2018, barley and chickpea consumed more water due to more evaporation and higher mean temperature, especially in April to May (Table 2).

### Pigment contents

During the both years, irrigation regime and cropping system had noticeable effect on pigment contents of barley (Table 5). Early sowing of barley and chickpea in December (b_1_c_1_) in 2017, increased chlorophyll *a* content of barley as well as 2018. Under water stress, chlorophyll *a* content of barley was affected by water deficit in the late season, negatively and the highest amount was obtained in b_2_ and b_1_c_2_ treatments. The b_1_c_1_ and b_1_c_2_ intercropping treatments influenced chlorophyll *b* content of barley which in rage of 0.39 to 0.49 mg. g^-1^ FW in normal irrigation and 0.23 to 0.27 mg. g^-1^ FW under water stress (Table 5). Similar to chlorophyll *a*, water stress and late sowing of barley had significant effect on declining the chlorophyll *b* content of barley (0.07 to 0.11 mg. g^-1^ FW). The highest carotenoid contend was created by b_1_c_2_ under normal irrigating while in water stress conditions, b_2_c_2_ treatment in 2017 had the maximum carotenoid content (0.41 mg. g^-1^ FW) by no significant difference with b_1_c_2_, b_2_c_1_ intercropping treatments. In addition, early sowing of barley (b_1_c_1_ and b_1_c_2_) had better performance in terms of total chlorophyll compared to the late sowing of barley (b_2_c_1_ and b_2_c_2_) treatments. Increasing the growing season length in early barley cultivation seems to contribute to better establishment and adaptation of barley intercropped with chickpea, so that total chlorophyll content enhanced more than late barley cultivation (Table 5).

**Table 5.**
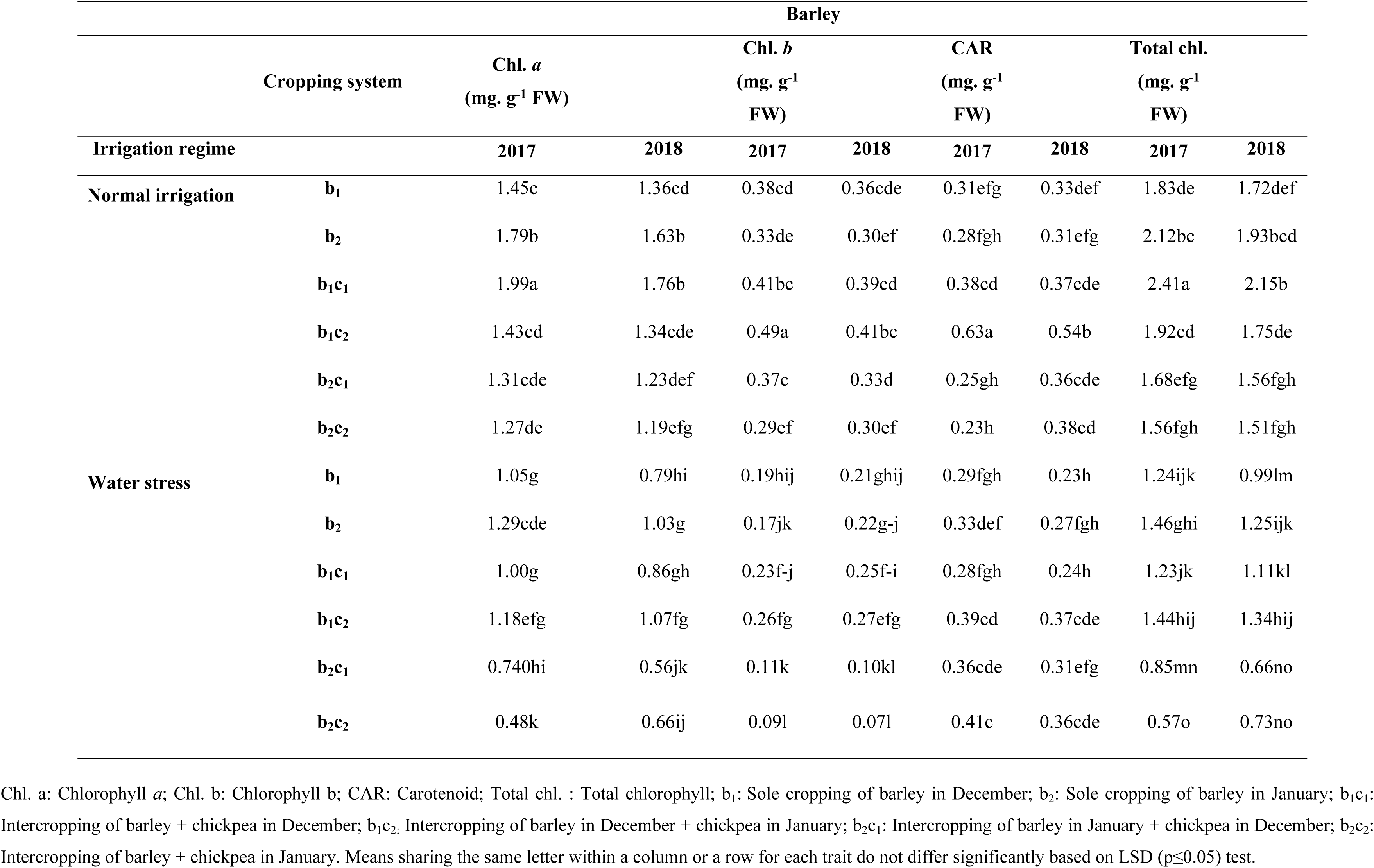
Interaction effect of irrigation regime and cropping system of barley with chickpea on chlorophyll *a*, *b*, carotenoid and total chlorophyll of barley.

In chickpea, pigment contents were affected by irrigation regime and cropping system during the two years of the experiment (Table 6). Late sowing of chickpea in January (b_1_c_2_ and b_2_c_2)_ enhanced chlorophyll *a* content of chickpea which in 2017 was more than 2018, significantly (p ≤ 0.05). Under water stress, chlorophyll *a* content in late sowing of chickpea with barley (b_2_c_2_) increased 25 and 29% compared to sole cropping of chickpea in January (c_2)_ in 2017 and 2018, respectively. In normal irrigation, the highest chlorophyll *b* content of chickpea was obtained in b_1_c_1_ (0.41 mg. mg. g^-1^ FW) and b_2_c_2_ (0.39 mg. mg. g^-1^ FW) treatments of the first year which was more than the second year. Similar trend was observed under water stress conditions and chlorophyll *b* content was in range of 0.21 to 0.31 mg. mg. g^-1^ FW (Table 6). During the 2017 and 2018, carotenoid content of chickpea was affected by cropping system so that, in both of the irrigation regimes, b_1_c_2_ and b_2_c_2_ had carotenoid content in range of 0.19 to 0.26 mg. g^-1^ FW in normal irrigation and 0.17 to 0.24 mg. g^-1^ FW under water stress (Table 6). In normal irrigation, b_2_c_1_ treatment and in water stress condition b_1_c_1_ and b_2_c_1_ treatments had the lowest carotenoid content of chickpea compared to the other intercropping treatments. It is appeared that late sowing of chickpea in January improved the carotenoid content of chickpea in comparison to early sowing in December. Finally, in intercropping treatments, total chlorophyll in b_1_c_2_ and b_2_c_2_ in normal irrigation and b_2_c_2_ in water stress condition enhanced compared to early sowing of chickpea intercropped with barley, significantly (p ≤ 0.05). Also, in both of the irrigation regimes, total chlorophyll in c_1_ and c_2_ treatments of the first year, significantly (p ≤ 0.05) was more than the second year. Under water shortage, when the growing season length of chickpea decreased in b_1_c_2_ and b_2_c_2_, due to better adaptation of chickpea in understory of barley rows, the total chlorophyll of chickpea enhanced compared to first sowing date (Table 6).

**Table 6.** Interaction effect of irrigation regime and cropping system of barley with chickpea on chlorophyll *a*, *b*, carotenoid and total chlorophyll of chickpea.

|  |  | Chickpea |  |  |  |  |  |  |  |
| --- | --- | --- | --- | --- | --- | --- | --- | --- | --- |
| Irrigation regime | Cropping system | Chl. *a*<br>(mg. g<sup>-1</sup> FW) |  | Chl. *b*<br>(mg. g<sup>-1</sup> FW) |  | CAR<br>(mg. g<sup>-1</sup> FW) |  | Total chl.<br>(mg. g<sup>-1</sup> FW) |  |
|  |  | 2017 | 2018 | 2017 | 2018 | 2017 | 2018 | 2017 | 2018 |
| Normal Irrigation | c<sub>1</sub> | 1.25c | 1.11e | 0.33bc | 0.29d-g | 0.23abc | 0.22abc | 1.58bc | 1.40d |
|  | c<sub>2</sub> | 1.27bc | 1.09ef | 0.27d-h | 0.28d-h | 0.16de | 0.20bcd | 1.54bc | 1.37de |
|  | b<sub>1</sub>c<sub>1</sub> | 0.79h | 0.72hi | 0.41a | 0.32cde | 0.18cd | 0.17d | 1.20efg | 1.04hi |
|  | b<sub>1</sub>c<sub>2</sub> | 1.35b | 1.24c | 0.33bc | 0.30def | 0.22abc | 0.19bcd | 1.68b | 1.54bc |
|  | b<sub>2</sub>c<sub>1</sub> | 0.78h | 0.63j | 0.30def | 0.25e-h | 0.09f | 0.11ef | 1.08gh | 0.88jkl |
|  | b<sub>2</sub>c<sub>2</sub> | 1.48a | 1.22cd | 0.39abc | 0.33bc | 0.23abc | 0.26a | 1.87a | 1.55bc |
| Water stress | c<sub>1</sub> | 0.80h | 0.63j | 0.29d-h | 0.21hij | 0.22abc | 0.18cd | 1.09g | 0.84j |
|  | c<sub>2</sub> | 0.92g | 0.78h | 0.30def | 0.19g-j | 0.26a | 0.21a-d | 1.32def | 0.97hij |
|  | b<sub>1</sub>c<sub>1</sub> | 0.52k | 0.50k | 0.27def | 0.23ghi | 0.11ef | 0.09f | 0.79klm | 0.73lmn |
|  | b<sub>1</sub>c<sub>2</sub> | 0.72h | 0.68ij | 0.21hij | 0.24f-i | 0.19bc | 0.17d | 0.93ijk | 0.92ij |
|  | b<sub>2</sub>c<sub>1</sub> | 0.53k | 0.46k | 0.17ij | 0.15j | 0.11ef | 0.08f | 0.72mn | 0.61n |
|  | b<sub>2</sub>c<sub>2</sub> | 1.15de | 1.01f | 0.31def | 0.27def | 0.24ab | 0.21a-d | 1.46cd | 1.28e |

Chl. *a*: Chlorophyll *a*; Chl. *b*: Chlorophyll *b*; CAR: Carotenoid; Total chl. : Total chlorophyll; c<sub>1</sub>: Sole cropping of chickpea in December; c<sub>2</sub>: Sole cropping of chickpea in January; b<sub>1</sub>c<sub>1</sub>: Intercropping of barley + chickpea in December; b<sub>1</sub>c<sub>2</sub>: Intercropping of barley in December + chickpea in January; b<sub>2</sub>c<sub>1</sub>: Intercropping of barley in January + chickpea in December; b<sub>2</sub>c<sub>2</sub>: Intercropping of barley + chickpea in January. Means sharing the same letter within a column or a row for each trait do not differ significantly based on LSD (p≤0.05) test.

### Antioxidant enzymes Activity

The mean comparison data of catalase (CAT) and peroxidase (POX) activity of barley intercropped with chickpea under different irrigation regime during in 2017 and 2018 are presented in Table 7. Water stress affected the CAT activity of barley, positively. The highest amount of CAT was created by b_2_c_1_ treatment and in the second year was more than the first year. In the second year, because of higher temperature and evaporation during the reproductive stages (Table 2), barley might be responded to water stress by enhancing the CAT activity. On the other hand, intercropping of barley with chickpea improved the activity of CAT compared to sole cropping of barley (b_1_ and b_2_) when plants subjected to water stress. In normal irrigation, late sowing of barley in January (b_2_) declined the POX activity of barley compared to b_1_ treatment. In contrast, under water stress POX activity of b_2_ from 9.88 and 10.11 Units. mg^-1^ protein in 2017 and 2018, declined to 5.27 and 6.53 Units. mg^-1^ protein in b_1_, respectively (Table 7). In response to late sowing date of barley in January and escape to water deficit in the late season, POX activity increased in b_2_c_2_. The CAT activity of chickpea was promoted by water stress, so that by relay intercropping of chickpea in January (b_1_c_2_) reach to 5.22 and 4.31 Units. mg^-1^ protein in 2017 and 2018, respectively (Table 8). In two irrigation regimes, late sowing of chickpea in sole cropping (c_2_) had the lower CAT activity in range of 0.69 to 0.94 Units. mg^-1^ protein compared to c_1_. On the other hand, in intercropping treatments, b_1_c_1_ had the least CAT activity in range of 0.43 to 1.47 Units. mg^-1^ protein. The POX activity was influenced by water stress, positively and in b_1_c_2_ of normal irrigation from 2.13 and 2.98 Units. mg^-1^ protein enhanced to 11.43 and 11.9 Units. mg^-1^ protein in 2017 and 2018 respectively (Table 8). Likewise, in b_1_c_2_ POX activity of chickpea increased 141 and 86% compared to c_2_, in the first and second year, respectively. Under water stress, the late sowing date of chickpea in January intercropped with barley in December (b_1_c_2_) could be alleviate the water stress effect through increasing CAT and POX activities of chickpea.

**Table 7.** Interaction effect of irrigation regime and cropping system of barley with chickpea on catalase and peroxidase activity of barley.

|  |  | CAT<br>(Units. mg<sup>-1</sup><br>protein) |  | POX<br>(Units. mg<sup>-1</sup><br>protein) |  |
| --- | --- | --- | --- | --- | --- |
| Irrigation regime |  | 2017 | 2018 | 2017 | 2018 |
| Normal irrigation | $b_1$ | 0.40j | 0.45j | 4.7ghi | 4.81ghi |
| | $b_2$ | 0.64i | 0.67i | 1.30j | 1.39j |
| | $b_1c_1$ | 1.48g | 1.56fg | 4.11i | 4.21hi |
| | $b_1c_2$ | 0.44j | 0.65i | 4.73ghi | 4.73ghi |
| | $b_2c_1$ | 2.24c | 2.36ab | 4.02i | 4.91ghi |
| | $b_2c_2$ | 0.51j | 0.61i | 4.33ghi | 5.12ghi |
| Water stress | $b_1$ | 0.85h | 0.93h | 5.27gh | 6.53f |
| | $b_2$ | 1.48g | 1.56fg | 9.88bc | 10.11b |
| | $b_1c_1$ | 1.70e | 1.82d | 8.26de | 8.91cd |
| | $b_1c_2$ | 1.67ef | 1.77de | 5.41g | 7.23ef |
| | $b_2c_1$ | 2.26bc | 2.43a | 9.45c | 9.56c |
| | $b_2c_2$ | 1.57fg | 2.15c | 12.81a | 13.21a |

CAT: Catalase; POX: Peroxidase; $b_1$ : Sole cropping of barley in December; $b_2$ : Sole cropping of barley in January; $b_1c_1$ : Intercropping of barley + chickpea in December; $b_1c_2$ : Intercropping of barley in December + chickpea in January; $b_2c_1$ : Intercropping of barley in January + chickpea in December; $b_2c_2$ : Intercropping of barley + chickpea in January. Means sharing the same letter within a column or a row for each trait do not differ significantly based on LSD ( $p \leq 0.05$ ) test.

**Table 8.** Interaction effect of irrigation regime and cropping system of barley with chickpea on catalase and peroxidase activity of chickpea.

|  |  | CAT<br>(Units. mg<sup>-1</sup><br>protein) |  | POX<br>(Units. mg<sup>-1</sup> protein) |  |
| --- | --- | --- | --- | --- | --- |
| Irrigation regime |  | 2017 | 2018 | 2017 | 2018 |
| Normal irrigation | c<sub>1</sub> | 1.34ef | 1.11gh | 1.89mn | 1.61n |
|  | c<sub>2</sub> | 0.74ij | 0.69j | 3.33k | 2.53lm |
|  | b<sub>1</sub>c<sub>1</sub> | 0.45k | 0.43k | 3.43k | 3.03kl |
|  | b<sub>1</sub>c<sub>2</sub> | 1.35ef | 1.21fg | 2.13mn | 2.98k |
|  | b<sub>2</sub>c<sub>1</sub> | 0.86ij | 0.75i | 4.57j | 3.26k |
|  | b<sub>2</sub>c<sub>2</sub> | 1.21fg | 1.11gh | 3.32k | 3.43k |
| Water stress | c<sub>1</sub> | 1.48e | 1.36ef | 5.32hi | 6.28fg |
|  | c<sub>2</sub> | 0.94hi | 0.85ij | 4.74ij | 6.37fg |
|  | b<sub>1</sub>c<sub>1</sub> | 1.47e | 1.23f | 5.68gh | 6.27fg |
|  | b<sub>1</sub>c<sub>2</sub> | 5.22a | 4.31b | 11.43ab | 11.9a |
|  | b<sub>2</sub>c<sub>1</sub> | 2.47c | 2.15d | 6.91ef | 8.21d |
|  | b<sub>2</sub>c<sub>2</sub> | 2.23d | 2.21d | 9.47c | 10.93b |

CAT: Catalase; POX: Peroxidase; c<sub>1</sub>: Sole cropping of chickpea in December; c<sub>2</sub>: Sole cropping of chickpea in January; b<sub>1</sub>c<sub>1</sub>: Intercropping of barley + chickpea in December; b<sub>1</sub>c<sub>2</sub>: Intercropping of barley in December + chickpea in January; b<sub>2</sub>c<sub>1</sub>: Intercropping of barley in January + chickpea in December; b<sub>2</sub>c<sub>2</sub>: Intercropping of barley + chickpea in January. Means sharing the same letter within a column or a row for each trait do not differ significantly based on LSD (p≤0.05) test.

### Relative water content (RWC)

In all of the cropping treatments, the RWCs of barley in 2017 growing season were higher than 2018 (Fig. 2a). The higher rainfall, and lower mean temperature and evaporation in 2017 compared to 2018 (Table 2) could explain the seasonal differences in RWC. Also, in each intercropping treatments, RWCs in normal irrigation were more than water stress conditions, significantly (p ≤ 0.05). The highest RWCs of barley were obtained in b_1_c_2_, b_1_c_1_ and b_2_c_1_ intercropping treatments under normal irrigation regime by significant differences with barley sole cropping. Similarly, under water stress, barley intercropped with chickpea enhanced RWCs amount compared to barley sole cropping. On the other hand, the lowest RWC was observed in b_2_c_2_ with significant difference by the other intercropping treatments. The mean comparison data of RWC of chickpea under different cropping treatments and irrigation regimes during the 2017 and 2018 growing seasons are given in Fig. 2b. The highest RWCs of chickpea were obtained in relay intercropping treatments (b_2_c_1_ and b_1_c_2_) which had significant difference with sole cropping of chickpea. On the other hand, water stress influenced RWC in all of the cropping systems, negatively. It seems that simultaneous sowing of chickpea with barley in January (b_2_c_2_) could not improve the acclimation of chickpea to water stress because of shortening the growing season length and sensitivity of chickpea.

**Figure 2.**
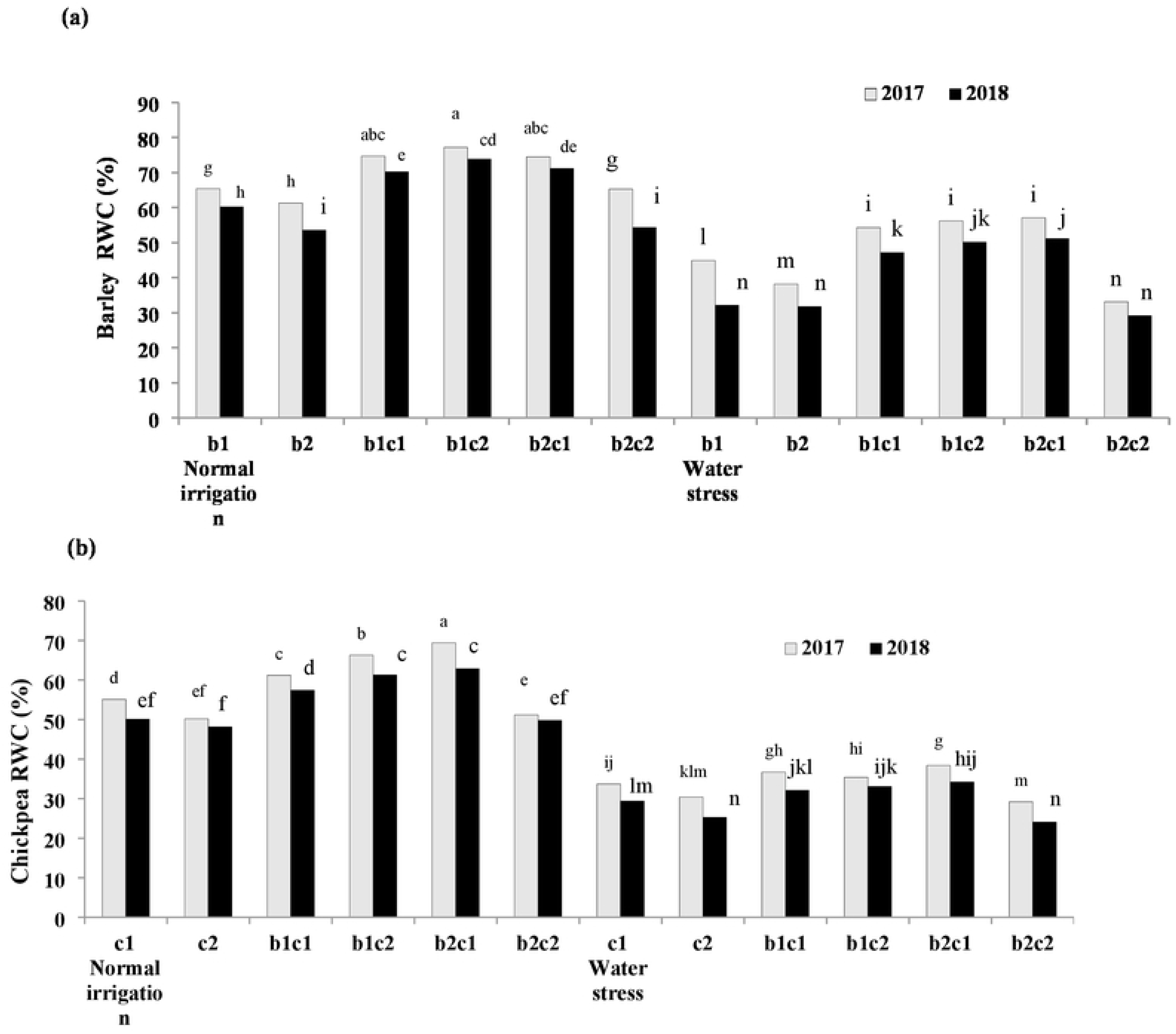
Interaction effect of irrigation regime and cropping system of barley with chickpea on relative water content (RWC) of barley (a) and chickpea (b). b_1_: Sole cropping of barley in December b_2_: Sole cropping of barley ion January, c_1_:Sole cropping of chickpea in December c_2_: Sole cropping of chickpea in January, b_1_c_1_: Intercropping of barley + chickpea in December; b_1_c_1_: Intercropping of barley 1n December + chickpea 1n January, b_2_c_1_: Intercropping of barley in January + chickpea in December, b_2_ci: lntercropping of barley + chickpea in January. Means sharing the same letter do not differ significantly based on LSD <u>(p<0.05)</u> test.

### Grain yield of barley and chickpea

The interaction effect of irrigation regime and intercropping treatments of barley with chickpea on grain yield of barley are shown in Fig. 3a. Under normal irrigation, relay intercropping of chickpea with barley in b_1_c_2_, increased barley grain yield by 17 and 18% compared to sole cropping of barley in December (b_1_). Water stress, affected the grain yield of barley, negatively, however in b_1_c_2_ treatment grain yield of barley enhanced 19 and 35% compared to b_1_. In contrast, in another relay intercropping treatment (b_2_c_1_), late sowing of barley could not mitigate adverse effects of water deficit on barley grain yield. Overall, barley grain yield in the first year was more than second year and in both irrigation regimes. In spite of barley, in normal irrigation regime, sole cropping of chickpea in December and January (c_1_ and c_2_) had the highest gran yield in range of 1721 to 2033 kg ha^-1^. Under water stress, the grain yield of chickpea in b_2_c_1_ was higher than c_1_ treatment with no significant difference (Fig. 3b). Early sowing of barley in December intercropped with late sowing of chickpea in January improved its competition ability to alleviate the negative effect of barley by faster growth rate. The lower chickpea grain yield in simultaneous sowing of barley and chickpea in January (b_2_c_2)_ might be related to shortening the growth period of chickpea and less ability to compete with barley under water stress.

**Figure 3.**
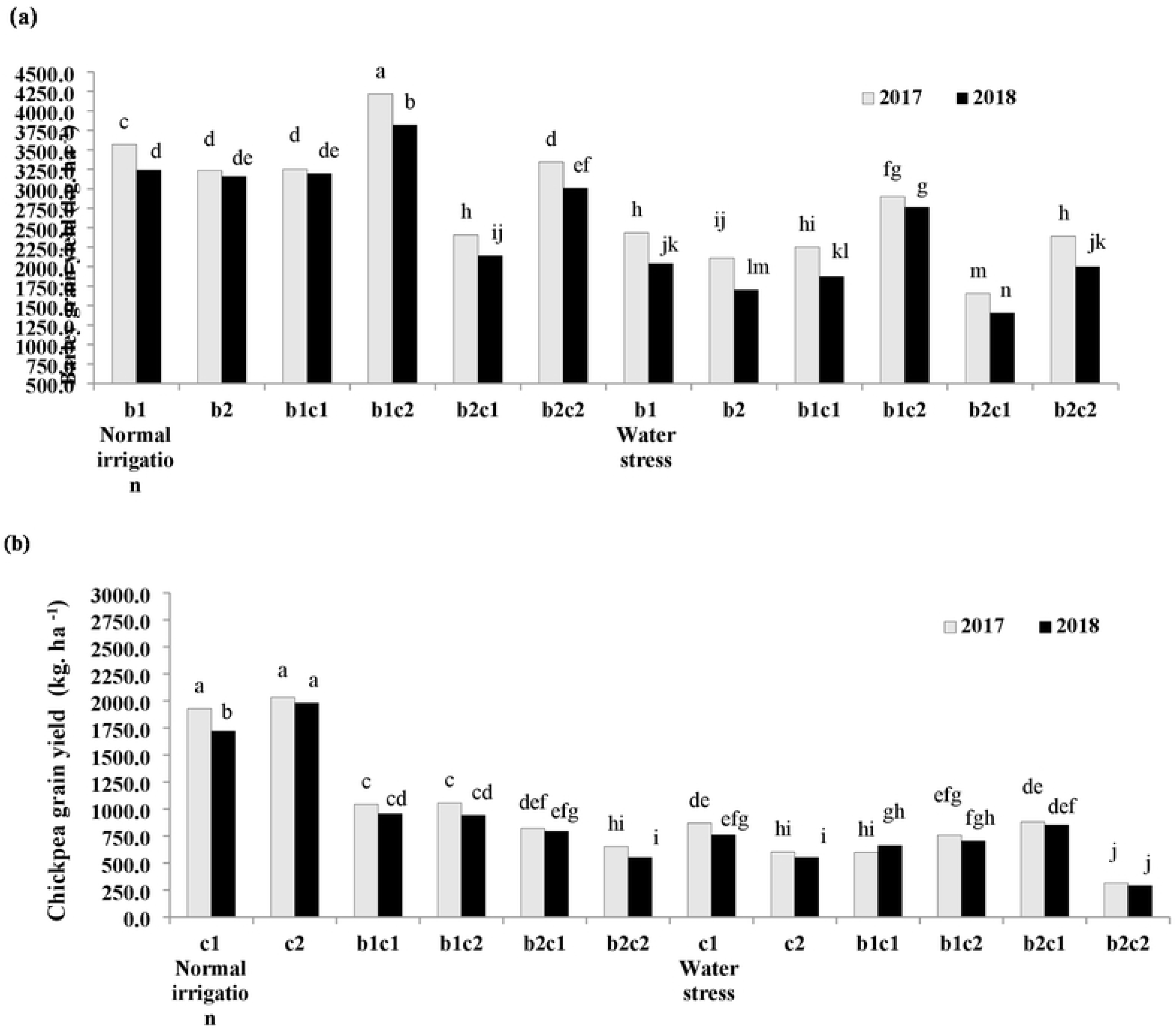
Interaction effect of irrigation regime and cropping system of barley with chickpea on grain yield of barley (a) and chickpea (b). b_1_: Sole cropping ofbarley in December b_2_: Sole cropping of barley in January, C_1_: Sole cropping of chickpea in December c2: Sole cropping of chickpea m January, b_1_c_1_: lntercropping of barley + chickpea m December; b_1_c_1_, lntercropping of barley in December + chickpea in January, b_2_c_1_: lntercropping of barley in January + chickpea in December, b_2_c_1_: Intercropping of barley + chickpea in January. Means sharing the same letter do not differ significantly based on LSD (p<O.OS) test.

### Biological yield of barley and chickpea

The biological yield of barley was influenced by irrigation regime and cropping system, so that the higher amount was observed in 2017 compared to 2018 (Fig. 4a). The b_1_c_2_ intercropping treatment with the maximum biological yield had the best intercropping treatment by 19 and 14% increase as compared to b_1_ in 2017 and 2018, respectively. Similar trend was observed in water stress conditions so that biological yield from 5244 and 4902 kg ha^-1^ reached to 6212 and 5986 kg ha^-1^ in b_1_c_2_ treatment during 2017 and 2018, respectively. The b_2_c_1_ treatment with 3701 kg ha^-1^ in 2017 and 3501 kg ha^-1^ under normal irrigation had not acceptable biological yield compared to the other treatments. In chickpea similar to grain yield, in normal irrigation sole cropping increased the biological yield in range of 4982 to 5466 kg ha^-1^ due to no competition with barley (Fig 4b). Controversy, under water stress, the amounts of biological yields in intercropping were close to sole cropping except the b_2_c_2_ treatment. Late sowing of chickpea with barley in January (b_2_c_2_), enhanced interspecific competition which declined biological yield of chickpea more than barley especially under water stress (Fig 4b).

**Figure 4.**
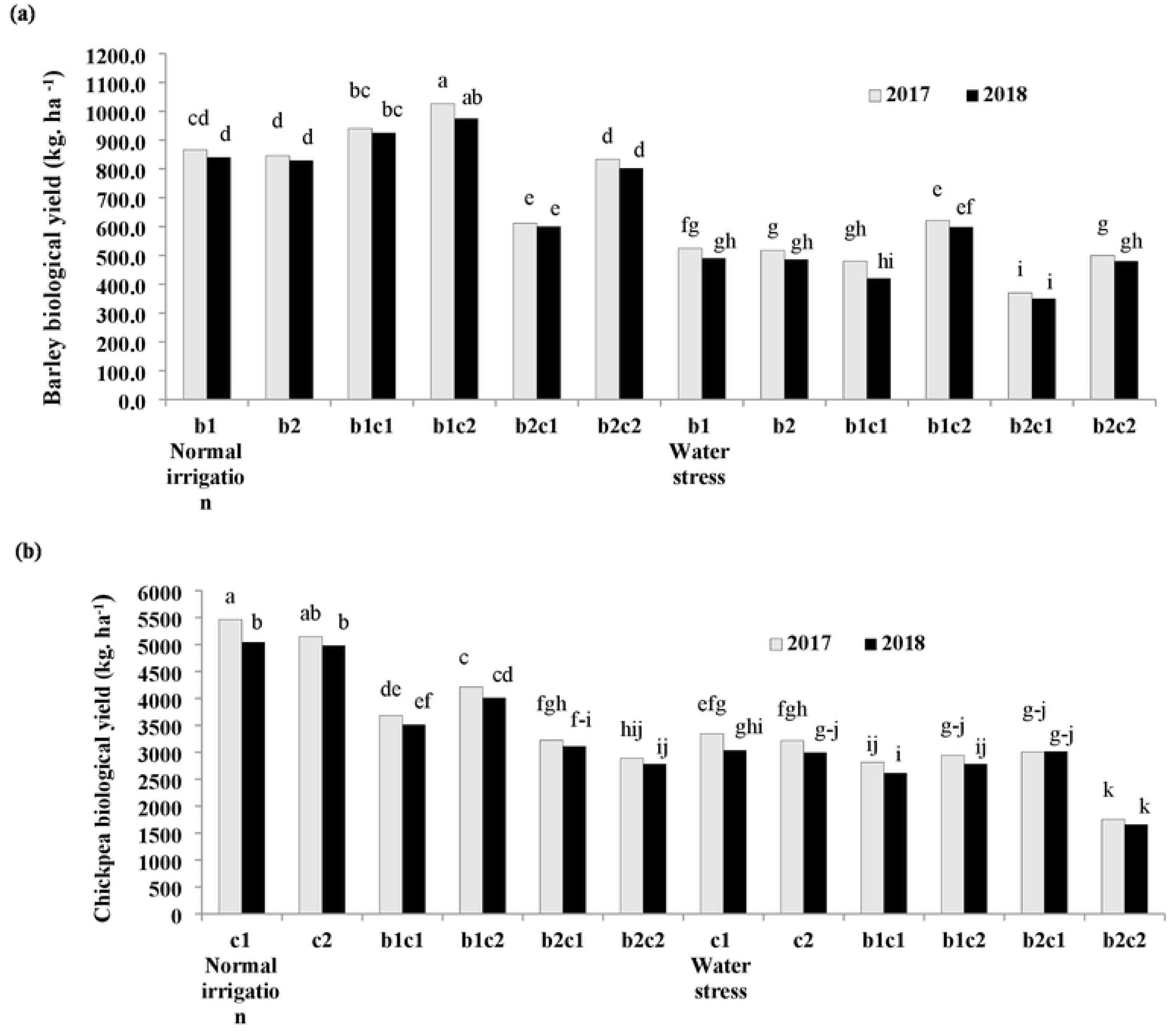
Interaction effect of irrigation regime and cropping system of barley with chickpea on biological yield of barley (a) and chickpea (b). bi: Sole cropping of barley in December b2: Sole cropping of barley in January, c_1_: Sole cropping of chickpea in December c_2_: Sole cropping of chickpea in January, b_1_c_1_: Intercropping of barley+ chickpea in December; b_1_ei: Intercropping of barley in December+ chickpea in January, b_2_c_1_: Intercropping of barley in January + chickpea in December, b_2_c_2_: lntercropping of barley + chickpea in January. Means sharing the same letter do not differ significantly based on LSD <u>(p<0.05)</u> test.

### Harvest index of barley and chickpea

In each year, intercropping of barley with chickpea under water stress affected the harvest index (HI) of barley in comparison to b_2_ treatment, significantly (Fig. 5a). Overall water stress increased HI of barley and in 2017 was more than 2018. This is likely due to differences in growing season climatic conditions as previously discussed (Table 2). In normal irrigation, late sowing of chickpea in January (c_2_), enhanced the HI by 39.5% and 39.8% in 2017 and 2018, respectively (Fig. 5b). In contrast, under water stress, HI in c_2_ declined sharply as compared to b_1_c_1_, b_1_c_2_ and b_2_c_1_ treatments. Declining of HI in C_2_ treatment under stress might be attributed to shortening the growth period and increasing the intraspecific competition for water in the late season. The HI of chickpea in relay intercropping of barley in January with chickpea in December (b_2_c_1_) enhanced 29.3% in 2017 and 28.3% in 2018. under water stress.

**Figure 5.**
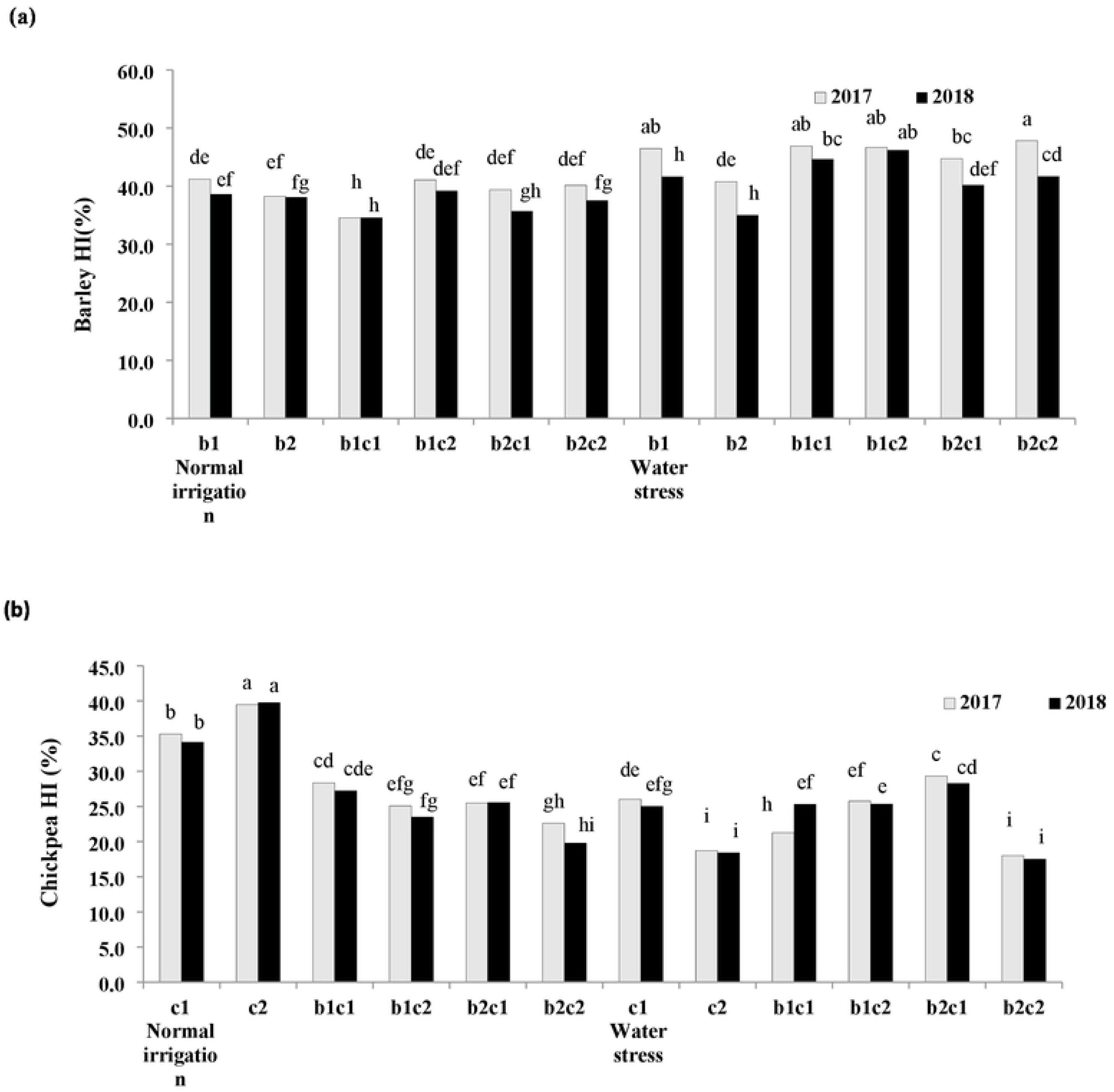
Interaction effect of irrigation regime and cropping system of barley, vith chickpea on harvest index (HI) of barley (a) and chickpea (b). b_1_: Sole cropping of barley in December b_2_: Sole cropping of barley in January, c_1_: Sole cropping of chickpea in December c_2_: Sole cropping of chickpea in January, b_1_c_1_: lntercropping of barley + chickpea in December; b_1_c_2_, Intercropping of barley in December+ chickpea in January, b_2_c_1_: lntercropping of barley in January + chickpea in December, b_2_ei: lntercropping of barley+ chickpea in January. Means sharing the same letter do not differ significantly based on LSD (pg).05) test.

### Water use efficiency

The water use efficiency (WUE) in each irrigation regime and cropping system in 2017 and 2018 are presented in Fig. 6. Results showed that in all of the treatments WUE in 2017 was higher than 2018. The highest WUE was obtained in b_1_c_2_ under both of the normal irrigation and water stress in 2017 growing season. On the other hand, WUE in c_2_ treatment was less than the other sole cropping treatments. Early sowing of barley in December had significant effect on WUE as compared to late sowing date in January (b_2_). The increasing the WUE in b_1_c_2_ might be related to more acclimation of barley and chickpea to environmental conditions, especially under late season water stress. Also, the higher rainfall and lower mean temperature and evaporation in the first year could be caused more WUE than the second year.

**Figure 6.**
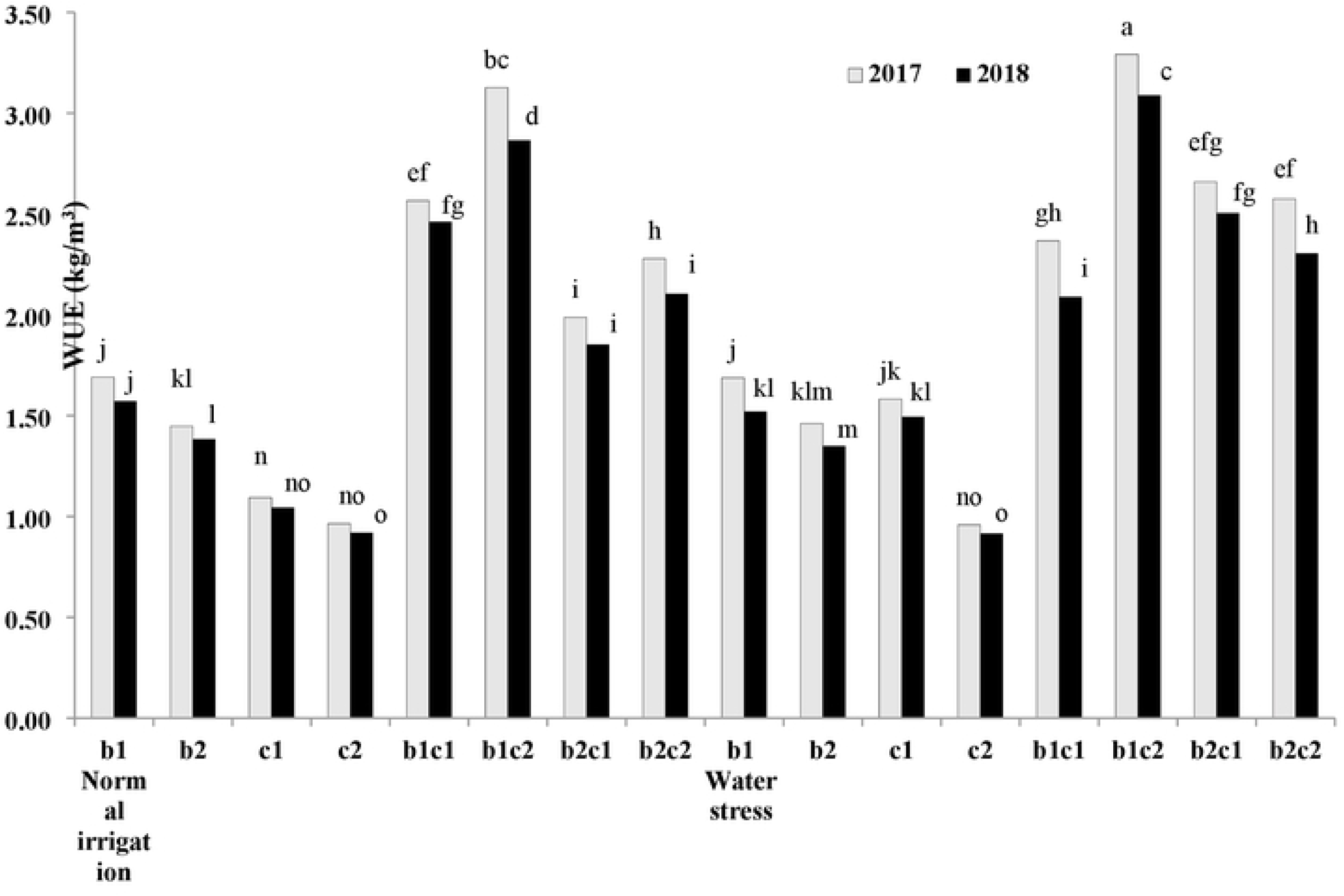
Interaction effect of irrigation regime and cropping system of barley with chickpea on water use efficiency (WUE) of barley (a) and chickpea (b). bi: Sole cropping of barley in December b_2_: Sole cropping of barley in January, c_1_: Sole cropping of chickpea in December ei: Sole cropping of chickpea in January, b_1_c_1_: lntercropping of barley+ chickpea in December; b_1_c_2_, Intercropping of barley in December + chickpea in January, b_2_c_1_: Intercropping of barley in January + chickpea in December, b2ei: Intercropping of barley + chickpea in January. Means sharing the same letter do not differ significantly based on LSD <u>(p<0.05)</u> test.

### Land equivalent ratio

In both of the irrigation regimes, LER of barley (LER_b_) in b_1_c_2_ treatments was in range of 1.15 to 1.18 which was more than the other intercropping treatments, significantly (p ≤ 0.05) (Table 9).

**Table 9.** Interaction effect of irrigation regime and cropping system on land equivalent ratio and competition ratio of barley with chickpea intercropping.

| Irrigation regime | Cropping system | LER |  |  |  |  |  | CR |  |  |  |  |  |
| --- | --- | --- | --- | --- | --- | --- | --- | --- | --- | --- | --- | --- | --- |
|  |  | Barley |  | Chickpea |  | Total |  | Barley |  | Chickpea |  | Total |  |
|  |  | 2017 | 2018 | 2017 | 2018 | 2017 | 2018 | 2017 | 2018 | 2017 | 2018 | 2017 | 2018 |
| Normal irrigation | b<sub>1</sub>c<sub>1</sub> | 1.09b | 1.10b | 0.67de | 0.70cd | 1.76bc | 1.80b | 1.61bcd | 1.58cd | 0.62fg | 0.63fg | 2.23c | 2.21c |
|  | b<sub>1</sub>c<sub>2</sub> | 1.18a | 1.16a | 0.82bc | 0.81bc | 2.00a | 1.97a | 1.45de | 1.44de | 0.69ef | 0.68ef | 2.14d | 2.12d |
|  | b<sub>2</sub>c<sub>1</sub> | 0.71f | 0.72f | 0.59def | 0.62def | 1.30g | 1.34fg | 1.20fg | 1.17fg | 0.83cd | 0.84cd | 2.03f | 2.01f |
|  | b<sub>2</sub>c<sub>2</sub> | 0.98c | 0.97c | 0.56ef | 0.54f | 1.55de | 1.51ef | 1.76abc | 1.79ab | 0.57g | 0.56g | 2.33b | 2.35b |
| Water stress | b<sub>1</sub>c<sub>1</sub> | 0.92d | 0.86e | 0.84b | 0.86b | 1.76bc | 1.72bcd | 1.09fg | 1.01g | 0.92bc | 1.02b | 2.01f | 2.03f |
|  | b<sub>1</sub>c<sub>2</sub> | 1.18a | 1.15a | 0.92ab | 0.93ab | 2.10a | 2.08a | 1.29ef | 1.25ef | 0.77de | 0.80d | 2.07e | 2.05ef |
|  | b<sub>2</sub>c<sub>1</sub> | 0.72f | 0.70f | 0.90ab | 0.99a | 1.61cde | 1.69bcd | 0.80h | 0.71h | 1.26b | 1.42a | 2.05ef | 2.12d |
|  | b<sub>2</sub>c<sub>2</sub> | 0.97c | 0.99c | 0.55f | 0.53f | 1.51ef | 1.52def | 1.77abc | 1.86a | 0.56g | 0.54g | 2.34b | 2.40a |

LER: Land equivalent ratio; CR: Competition ratio; b<sub>1</sub>c<sub>1</sub>: Intercropping of barley + chickpea in December; b<sub>1</sub>c<sub>2</sub>: Intercropping of barley in December + chickpea in January; b<sub>2</sub>c<sub>1</sub>: Intercropping of barley in January + chickpea in December; b<sub>2</sub>c<sub>2</sub>: Intercropping of barley + chickpea in January. Means sharing the same letter within a column or a row for each trait do not differ significantly based on LSD (p≤0.05) test.

The LER of chickpea (LER_c_) in all of the intercropping treatments was higher than 0.5 where demonstrated the advantage of chickpea intercropped with barley as compared to sole cropping of chickpea (Table 9). LER total (LER_t_) was affected by irrigation regime and cropping system, so that in both of the irrigation regime, in b_1_c_2_ maximized, significantly (p≤ 0.05). In spite of the other treatment, in b_2_c_1_ under water stress LER_t_ increased significantly in comparison to normal irrigation (Table 9). In each intercropping treatment, there were no significant difference between the LER_t_ of 2017 and 2018 growing seasons as well as LER_b_ and LER_c_. Also, in all of the treatment, the LER_b_ was morer than LER_c_ which described the higher efficiency of barley in absorbing the water and light compared to chickpea.

### Competition ratio

The competition ratio (CR) was influenced by irrigation regime and intercropping treatment, significantly (p ≤ 0.05) (Table 9). In b_2_c_2_ treatment of two irrigation regimes, CR of barley (CR_b_) was more than the other intercropping treatment in range of 1.76 to 1.86. In b_2_c_1_ treatment under water stress, CR_b_ reached to 0.80 in 2017 and 0.71 in 2018 by significant differences (p ≤ 0.05) with the other treatments. Water stress increased the CR of chickpea (CR_c_) in b_1_c_1_ and b_2_c_1_,

significantly while in b_2_c_2_ declined to 0.54 to 0.57 under two irrigation regimes (Table 9). Late sowing of barley with chickpea simultaneously (b_2_c_2_) in both of the irrigation regimes, enhanced the CR total (CR_t_) compared to the other treatment (Table 9). In each intercropping treatment, there were no significant differences (p ≤ 0.05) between CR_t_ of 2017 and 2018 growing seasons.

## Discussion

In arid areas, water stress is threating agricultural sustainability, and strip-intercropping may serve as a suitable approach to mitigate the challenge. In south of Iran by increasing the temperature there was no sufficient rainfall during the reproductive stage of crop when the water requirement of barley and chickpea enhanced. Because of occurrence of rain fall in the cool season, the farmers prefer to culture the crops in December however, irrigated the crop in the warm season [46, 9]. In similar study on barley- field bean (*Vicia faba* L.) intercropping, Pampana et al. [47] found that the different rainfall amount of the two growing seasons influenced barley sole crop. Strip-intercropping increases the spatial distributions of soil water across the 0–110 cm rooting zones, improves the coordination of soil water sharing during the co-growth period, and provides compensatory effect for available soil water. The intercropped chickpea used soil water mostly in the top 20-cm layers, whereas cereals were able to absorb water from deeper-layers of the neighboring pea strips [19]. Chickpea extracted soil water mostly from the shallow (in the top 20 cm) soil depths and the majority of pea roots are concentrated in the 0–30 cm soil profile [33]. In our study, under water stress different root distribution might be declined total water applied in relay intercropping (b_1_c_2_ and b_2_c_1_) in comparison to sole cropping of barley (b_1_ and b_2_) and chickpea (c_1_ and c_2_) (Fig. 1).

Chlorophyll content has been a key parameter applied to evaluate the status of a crop especially under water stress [16, 26]. On the other hand, intercropping enhanced the chlorophyll content of leaf with increasing the nutrient and water availability [48]. Machiani et al. [49] reported that intercropping systems demonstrated higher chlorophyll contents compared to sole cropping. Maffei and Mucciarelli [50] declared that peppermint intercropped with soybean created higher chlorophyll and carotenoid contents and biological yield related to to sole cropping. Liu et al. [17] showed that chlorophyll *a* content was decreased sharply in peanut during water stress, while chlorophyll *b* content was approximately constant. In the present study, total chlorophyll of barley in all of the cropping treatments decreased significantly (p ≤ 0.05) when plants subjected to water stress. In barley, sowing of barley in December intercropped with chickpea in December and January (b_1_c_1_ and b_1_c_2_) created a suitable condition in pigment contents improvement (Table 5). In contrast, in chickpea seems that simultaneous sowing of barley and chickpea in January (b_2_c_2_) caused a favorite situation in increasing pigment content (Table 6). Singh and Aulakh [5] concluded that the more chlorophyll content in intercropping than sole wheat could be related to nitrogen transfer with chickpea and more soil moisture in wheat intercropped with chickpea.

When crops were exposed to severe water stress, the reactive oxygen species (ROS) like hydrogen peroxide and superoxide accumulated in the leaves. In this condition, crop enhances antioxidant contents of their leaves to alleviate the negative effects of ROSs [51]. Water stress retarded the photosynthetic abilities and growth rate of crop due to breakdown of the balance between the antioxidant contents such as catalase (CAT) and peroxidase (POX) and the production of ROSs [52]. Increase in CAT activity is a common response to water stress demonstrate prominent role of CAT in the leaf protection against chlorophyll oxidation [53]. Similar to our results, Mafakheri et al. [46] showed a higher CAT and POX activity under stress in three chickpea genotypes. Nair et al. [54] asserted that CAT and POX enhanced significantly in cowpea (*Vigna unguiculata* L.) when crop exposed to water stress. Little studies have been published in terms of intercropping effect on antioxidant activity of the crops. In one of the few studies, Eskandari and Alizadeh Amraie [9] reported that interaction effect of crop system and irrigation regime of Persian clover intercropped with wheat were significant (p≤ 0.05) on POX activity and the maximum activity of POX was observed under water stress which was 20% higher than wheat. In the present study, intercropping of barley with chickpea enhanced their capacity against to oxidation under water stress via increment the CAT and POX activities, significantly as compared to sole cropping of normal irrigation regime. In the current study, relay intercropping of chickpea in January with barley on December (b_1_c_2_) increased the enzymes activity of chickpea more than barley (Tables 7 and 8). This might be related to more sensitivity of chickpea to water deficit and less its competition ability in intercropped with barley [5, 35].

The RWC represents the water status of plant which is related to cell turgidity of the leaves. Also, cell division and development are closely related to cell turgidity, and positively affect RWC and grain yield [55, 56]. The tolerant plants maintain more water in their leaves because of more RWC as compared to sensitive plants [26]. Our findings are in agreement to Eskandari and Alizadeh Amraie [9] who declared that in all cropping systems, RWC of Persian clover and wheat declined by decreasing available water in the soil under water stress conditions. In our study, RWC in barley was more than chickpea in both of the irrigation regimes and cropping systems, which suggesting that barley was a more drought tolerant in comparison to chickpea. Also, RWC improved in barley and chickpea when intercropped together. It is appeared that intercropping barley with chickpea especially in b_1_c_2_ could be mitigated the adverse effects of water stress on crop growth rate by maintaining higher RWC in their leaves. Overall, the higher RWC of barley in b_1_c_1_ and b_1_c_2_ treatments could be created a better condition in total chlorophyll increment (Table 5).

In intercropping of cereals with legumes the yield of each crop could be affected by interspecific competition for crucial growth resources, allelopathic effects, water stress, sowing date and plant density of each crop caused in lower grain yield in intercrops [1, 5, 57]. Abu-Bakar et al. [11] declared that the highest grain yield of barley was obtained in sole crop compared to barley intercropped with lentil. In the current study, the lower chickpea grain yield in intercropping could be related to less competitive ability of chickpea in term of light, water and nutrient compared to sole cropping (Ahlawat et al. 2005). The higher production in sole cropping can be related to the homogeneous conditions under sole cropping [5]. Controversy, some researches showed that the grain yield and biological yield improved in intercropping by different crops and environments. For example, De la Fuente et al. [58] declared that sunflower and soybean intercropping created higher grain yield compared to sole cropping which could be related to complementary use of resource in space and time in intercropping systems. Also, Amossé et al. [59] declared that the wheat canopies had no inhibition effects on seedling establishment of legumes in relay intercropping. Känkänen and Eriksson [60] observed that legumes with intercropped barley had the lowest negative effect on barley yield compared to non-fixing annual relay intercrops because of slower growth of legumes in the early season. Galanopoulou et al. [31] reported that barley can be grown with faba bean due to crops created a high biological yield and grain yield by more exploiting the resources compared to monoculture. Recently, Luhmer et al. [61] reported that barley and poppy (*Papaver somniferum* L.) intercrops produced higher poppy yields compared to sole cropping, whereas early sowing dates of barley enhanced the its competition ability. Latati et al. [62] suggested that legumes facilitated the cereals production through optimum use of environmental resources in a suitable intercropping system. Iliadis (2001) declared that autumn winter sown chickpea created more grain yield than spring sown. In the current study, the early sowing of barley in December intercropped with chickpea in January (b1c2) increase the grain yield (Fig. 3a) and biological yield (Fig. 4a) of barley due to early suppression of chickpea by vigorous barley. The reduction chickpea yield in late sowing date of chickpea (December vs. January) might be due to declining in light transmission and interception in the lower canopy, leading to growth and development depression [34]. Under water stress, greater biological yield of barley in b_1_c_2_ treatments leading to higher grain yield, than sole cropping which possibly related to increase the total chlorophyll (Table 5) and (RWC) (Fig. 2a) and light interception by greater canopy at early sowing date of barley [64]. In contrast, late sowing date of barley intercropped with chickpea in b_2_c_1_ treatment enhanced chickpea grain yield. Delay in sowing of barley into the intercrop enhanced the suppression ability of chickpea over barley to gain the more growth resources. The early sowing date of the legume intercropped with cereals facilitates its seedling establishment, allowing to add more dry matter [59]. The simultaneous sowing date of chickpea with barley in January (b_2_c_2_) decreased the HI of chickpea compared to other intercropping treatments. The late sowing date of chickpea increased the shading of barley on chickpea which could be declined overall photosynthetic production to a level where chickpea compensated with decreasing the amount of assimilate allocated to grain filling and HI decreased, drastically. It is appeared that in b_1_c_2_ treatment, each crop occupied and accessed to growth resources from different ecological niches at different times due to delay intercropping while minimizing competitive interactions. Overall, grain yield, biological yield and HI of chickpea in b_2_c_2_ was less than b_1_c_1_ intercropping treatment because of shortening the growing season and met the grain filling period to late season water stress.

The higher LER in intercropping revealed the advantages of intercropping of cereals and legumes because of better utilization of resources like light, nutrients uptake and available water [29, 31, 65]. Galanopoulou et al. [31] declared that in all of intercrop treatment, the LERs of barley were higher than 0.5 but in faba bean were lower than 0.5, which demonstrated the advantages of barley compared to faba bean. Hauggaard-Nielsen et al. [66] reported that in pea intercropped with barley, the LER of chickpea decreased while partial LER of barley increased, significantly. In our study LER amounts of barley and chickpea were more than 0.5 while in all of the intercropping treatments the LER of barley was higher than chickpea. Similar to our results, Hamzei and Seyedi [67] declared that in all intercropping treatments of barley with chickpea the LER total was higher than one. It’s demonstrated the superiority of relay intercropping especially in b_1_c_2_ treatments compared to sole cropping. <u>Eskandari and Alizadeh</u> <u>Amraie</u> [9] in a similar study declared that LER of wheat- Persian clover intercropping under water stress was more than normal irrigation. They suggested that intercropping mitigated the adverse effects of water deficit by RWC and WUE increment of each crop resulting in LER increment. Chen et al. [19] reported that the corn-chickpea intercropping increased maize grain yield by an average (three-years) of 25% and enhanced maize WUE by 24%. In the dry areas with high soil evaporation, the increased systems productivity and WUE with intercropping is partly attributable to water sharing through possible water movement between the rooting zones and water compensation from one strip to the other. Fan et al. [68] reported that maize–pea intercropping produced 23–38% greater total yield than corresponding sole crops, as the LER ranged from 1.23 to 1.38 and intercropping improved water capture by plants compared with sole maize. In the current study, one of the reasons for increasing LER_t_ especially under water stress in intercropping of barley in December + chickpea in January (b_1_c_2_) was enhancing the grain yield (Fig. 3a), RWC (Fig 2a) and WUE (Gig 6) of barley compared to sole cropping.

CR reveals a useful evaluation of competition ability between two crops in intercropping. Machiani et al. [49] found that the CR amount of peppermint was more than 1 which was higher than soybean showing a yield superiority of peppermint intercropped with soybean. Andrade et al. [69] suggested that avoiding the overlapping of critical growth stages by relay intercropping improves resources use between intercrop components. They also found that yield advantage for intercrop is lower under scarce water availability and mainly associated with a decrease in intercropped legume productivity. Veisi et al. [13] declared that chickpea had no ability to compete with other plants because of its slow growth relatively in the seedling establishment stage. Greater competition ability of cereals in intercropping with legumes may be related to that cereals take up more water and nitrogen in the early season and accumulated more dry matter which caused shad on the legume and thereby reduce its competition ability in intercropping system [31, 66]. The more CR of barley compared to chickpea in all intercropping systems and irrigation regimes except the b_2_c_1_ treatments, was in consistent with the results of Megawer et al. [70]. In our study, the higher CR of barley compared to chickpea demonstrated the higher aggressively and superiority of barley and its capability in up taking of more resources compared with chickpea.

## Conclusions

Relay intercropping of barley-chickpea tended to be more productive as early sowing of barley in December intercropped with late sowing of chickpea in January (b_1_c_2_) in the south of Iran. In the second year, due to higher temperature and evaporation, barley- chickpea intercropping responded to water stress by enhancing enzymes activity more than first year. Under water stress, early sowing of barley had better performance in terms of total chlorophyll compared to the late sowing of barley. By decreasing growing season length of chickpea in b_1_c_2_ and b_2_c_2_ treatments, due to better adaptation of chickpea in understory of barley rows, the total chlorophyll of chickpea enhanced under water stress. In relay intercropping, using crops by contrasting rooting structure and suitable sowing date of each component, productivity may be improved through enhancing biochemical properties, relative water content and water use efficiency. It is concluded that relay intercropping of barley in December with chickpea in January could be a suitable intercropping system for sustainable agriculture under water stress.

## Acknowledgments

The authors thank to Abdullah Setodeh for his assistance in the lab, as well as the Agriculture and Natural Resources Research Center of Darab, Fars Province, Iran for providing the seeds of this research.

## Author Contributions

**Conceptualization:** Ehsan Bijanzadeh

**Data curation:** Negin Mohavieh Assadi

**Formal analysis:** Negin Mohavieh Assadi

**Funding acquisition:** Ehsan Bijanzadeh

**Investigation:** Negin Mohavieh Assadi

**Methodology:** Negin Mohavieh Assadi

**Project administration:** Negin Mohavieh Assadi, Ehsan Bijanzadeh

**Resources:** Negin Mohavieh Assadi

**Software:** Negin Mohavieh Assadi, Ehsan Bijanzadeh

**Supervision:** Ehsan Bijanzadeh

**Validation:** Negin Mohavieh Assadi, Ehsan Bijanzadeh

**Visualization:** Negin Mohavieh Assadi, Ehsan Bijanzadeh

**Writing – original draft:** Negin Mohavieh Assadi, Ehsan Bijanzadeh

**Writing – review & editing:** Ehsan Bijanzadeh

## References

1. Lithourgidis A, Vlachostergios D, Dordas C, Damalas C. Dry matter yield, nitrogen content, and competition in pea–cereal intercropping systems. Eur J Agr. 2011; 34: 287–294. https://doi.org/10.1016/j.eja.2011.02.007

2. Raza MA, Khalid MHB, Zhang X, Feng LY, Khan I, Hassan MJ, Ahmed M, Ansar M, Chen YK, Fan YF. Effect of planting patterns on yield, nutrient accumulation and distribution in maize and soybean under relay intercropping systems. Sci Rep. 2019; 9: 1–14. https://doi:10.1038/s41598-019-41364-1 PMID: 3089462.

3. 3. Ikram ul Haq M, Maqbool MM, Ali A, Farooq S, Khan S, Saddiq MS, Khan KA, Al, S, Ifnan Khan M, Hussain A. Optimizing planting geometry for barley-Egyptian clover intercropping system in semi-arid sub-tropical climate. Plos One. 2020; 15: e0233171. https://doi.org/10.1371/journal.pone.0233171

4. Inal A, Gunes A, Zhang F, Cakmak I. Peanut/maize intercropping induced changes in rhizosphere and nutrient concentrations in shoots. Plant Phy Bio. 2007; 45: 350–356. https://doi.10.1016/j.plaphy.2007.03.016 PMID: 17467283.

5. Singh B, Aulakh C. Effect on growth and yield of intercrops in wheat+ chickpea intercropping under limited nutrition and moisture. Indian J Eco. 2017; 44: 507–511.

6. Dhima K, Lithourgidis A, Vasilakoglou I, Dordas C. Competition indices of common vetch and cereal intercrops in two seeding ratio. Field Crops Res. 2007; 100: 249–256. https://doi.org/10.1016/j.fcr.2006.07.008

7. Coll L, Cerrudo A, Rizzalli R, Monzon J, Andrade FH. Capture and use of water and radiation in summer intercrops in the south-east Pampas of Argentina. Field Crops Res. 2012; 134:105–113. https://doi.org/10.1016/j.fcr.2012.05.005

8. Monzon JP, Mercau JL, Andrade J, Caviglia, OP, Cerrudo A, Cirilo AG, Vega CRC, Andrade FH, Calviño PA. Maize–soybean intensification alternatives for the Pampas.Field Cro Res. 2014; 162: 48–59. https://doi.org/10.1016/j.fcr.2014.03.012

9. Eskandari H, Alizadeh Amraie A. Antioxidant and yield responses of wheat and clover in intercropping system to late season water deficit induced by partial root-zone irrigation regime. J Plant Pro Fun. 2018; 6, 7–14.

10. 10. FAO., Food and agriculture organisation of the United Nations. 2019. Available from: http://faostat.fao.org

11. Abu-Bakar M, Riaz A, Ehsanullah Zahir AZ. Comparison of barley-based intercropping system for productivity and net economic return. Int J Agr Bio. 2014; 16: 1183–1188.

12. Chapagain T, Riseman A. Barley–pea intercropping: Effects on land productivity, carbon and nitrogen transformations. Field Crops Res. 2014; 166: 18–25. https://doi.org/10.1016/j.fcr.2014.06.014

13. Veisi M, Zand E, Moeini M M, Bassiri K. Review of research on weed management of chickpea in Iran: challenges, strategies and perspectives. J Plant Pro Res. 2020; 60: 113–125. https://doi.org/10.24425/jppr.2020.132212

14. Rady M, Gaballah MS. Improving barley yield grown under water stress conditions. Res J Rec Sci. 2012; 1: 1–6.

15. Amanullah Khalid S, Khalil F, Elshikh MS, Alwahibi MS, Alkahtani J. Growth and dry matter partitioning response in cereal-legume intercropping under full and limited irrigation regimes. Sci Rep. 2021; 11:12585. https://doi.org/10.1038/s41598-021-92022-4

16. Woo NS, Badger MR, Pogson BJ. A rapid, non-invasive procedure for quantitative assessment of drought survival using chlorophyll fluorescence. Plant Met. 2008; 4: 1–14. https://doi.org/10.1186/1746-4811-4-27

17. Liu X, Li L, Li M, Su L, Lian S, Zhang B, Li X, Ge K, Li L. AhGLK1 affects chlorophyll biosynthesis and photosynthesis in peanut leaves during recovery from drought. Sci Rep. 2018; 8: 2250. https://doi.org/10.1038/s41598-018-20542-7

18. Fang Y, Xiong L. General mechanisms of drought response and their application in drought resistance improvement in plants. Cel Mol Life Sci. 2015; 72: 673–689. https://doi.org/10.1007/s00018-014-1767-0 PMID: 25336153

19. Chen G, Kong X, Gan Y, Zhang R, Feng F, Yu A, Zhao C, Wan S, Chai Q. Enhancing the systems productivity and water use efficiency through coordinated soil water sharing and compensation in strip-intercropping. Sci Rep. 2018; 8:10494. https://doi.org/10.1038/s41598-018-28612-6

20. Li RH, Guo PG, Michael B, Stefania G, Salvatore C. Evaluation of chlorophyll content and fluorescence parameters as indicators of drought tolerance in barley. Agr Sci China. 2006; 5: 751–757. https://doi.org/10.1016/S1671-2927(06)60120-X

21. Miller G, Suzuki N, Ciftci-Yilmaz S, Mittler R. Reactive oxygen species homeostasis and signalling during drought and salinity stresses. Plant Cell Env. 2010; 33: 453–467. https://doi.org/10.1111/j.1365-3040.2009.02041.x

22. Zhang L, Zhang X, Wang K, Zhao Y, Zhai Y, Gao M, An Z, Liu J, Hu J. Foliar urea application affects nitric oxide burst and glycinebetaine metabolism in two maize cultivars under drought. Pak J Bot. 2012; 44: 81–86.

23. Gill SS, Tuteja N. Reactive oxygen species and antioxidant machinery in abiotic stress tolerance in crop plants. Plant Phy Bio. 2010; 48: 909–930. https://doi.org/10.1016/j.plaphy.2010.08.016

24. Sofo A, Scopa A, Nuzzaci M, Vitti A. Ascorbate peroxidase and catalase activities and their genetic regulation in plants subjected to drought and salinity stresses. Int J Mol Sci. 2015; 16: 13561–13578. https://doi.org/10.3390/ijms160613561 PMID: 26075872.

25. Laxa M, Liebthal M, Telman W, Chibani K, Dietz K J. The role of the plant antioxidant system in drought tolerance. Ant. 2019; 8: 94. https://doi.org/10.3390/antiox8040094 PMID: 30965652.

26. Bijanzadeh E, Emam Y, Pessarakli M. Biochemical responses of water-stressed triticale (*X Triticosecale wittmack*) to humic acid and jasmonic acid. J Plant Nut. 2021; 44: 252–269. https://doi.org/10.1080/01904167.2020.1806312

27. Yu Y, Stomph TJ, Makowski D, Vander Werf W. Temporal niche differentiation increases the land equivalent ratio of annual intercrops: a meta-analysis. Field Crops Res. 2015; 184: 133–144. http://dx.doi.org/10.1016/j.fcr.2015.09.010

28. Martin-Guay MO, Paquette A, Dupras J, Rivest D. The new green revolution: sustainable intensification of agriculture by intercropping. Sci Total Env. 2018; 615: 767–772. https://doi.org/10.1016/j.scitotenv.2017.10.024 PMID: 28992501

29. Lithourgidis A, Dordas C, Damalas CA, Vlachostergios D. Annual intercrops: an alternative pathway for sustainable agriculture. Aus J Crop Sci. 2011; 5: 396–410.

30. Bedoussac L, Journet EP, Hauggaard-Nielsen H, Naudin C, Corre-Hellou G, Jensen ES, Prieur L, Justes E. Ecological principles underlying the increase of productivity achieved by cereal-grain legume intercrops in organic farming. A review. Agr Sus Dev. 2015; 35: 911–935. https://doi.org/10.1007/s13593-014-0277-7

31. Galanopoulou K, Lithourgidis AS, Dordas CA. Intercropping of faba bean with barley at various spatial arrangements affects dry matter and N yield, nitrogen nutrition index, and interspecific competition. Not Bot Hor Agro Cluj-Nap. 2019; 47: 1116–1127. https://doi.org/10.15835/nbha47111520

32. Raza MA, Feng LY, Iqbal N, Ahmed M, Chen YK, Khalid MHB, Din AMU, Khan A, Ijaz W, Hussain A. Growth and development of soybean under changing light environments in relay intercropping system. PeerJ 2019; 7:e7262. https://doi.org/10.7717/peerj.7262

33. Wang Y, Qin Y, Chai Q, Feng F, Zhao C, Yu A. Interspecies interactions in relation to root distribution across the rooting profile in wheat-maize intercropping under different plant densities. Fro Plant Sci. 2018; 9: 1–17. https://doi.org/10.3389%2Ffpls.2018.00483 PMID: 29731758

34. Rahimi Azar M, Javanmard A, Shekari F, Pourmohammad A, Esfandyari E. Evaluation of yield and yield components chickpea (*Cicer arietinum* L.) in intercropping with spring barley (*Hordeum vulgare* L.). Cerc Agr Mold. 2013; 4: 75–85.

35. Ahlawat I, Gangaiah B, Singh O. Production potential of chickpea (*Cicer arietinum*) based intercropping systems under irrigated conditions. Indian J Agr. 2005; 50: 27–30.

36. Torkaman M, Mirshekari B, Farahvash F, Yarnia M, Ashraf Jafari A. Effect of sowing date and different intercropping patterns on yield and yield components of rapeseed (*Brassica napus* L.) and chickpea (*Cicer arietinum* L.). Leg Res. 2018; 41: 578–583.

37. Zadoks JC, Chang TT, Konzak CF. A decimal code for the growth stages of cereals. Weed Res. 1974; 14: 415–421. https://doi.org/10.1111/j.1365-3180.1974.tb01084.x

38. Ghazvini HO, Kohkan SA, Lakzadeh I, Fallahi HA, Alt Jafarbay J, Ghasemi M, Amini AA, Tabib Ghaffari SM, Sorkhi Lalelu B. Zehak, a new irrigated barley cultivar with wide adaptability in the warm and dry agro-climate zone in the south of Iran. Res Ach Field and Hort Cro. 2014; 3: 15–26.

39. Pirzahiri K, Kanouni H, Rokhzadi A. Response of some chickpea (*Cicer arietinum* L.) varieties to changes in plant density. J Crop Eco. 2020; 14: 293–310.

40. Arnon DI. Copper enzymes in isolated chloroplasts. Polyphenoloxidase in Beta vulgaris. Plant Phy. 1949; 24: 1–15. https://doi.org/10.1104%2Fpp.24.1.1

41. Aebi H. Catalase in vitro. Met Enz. 1984; 105: 121-126. https://doi.org/10.1016/s0076-6879(84)05016-3 PMID: 6727660

42. Chance B, Maehly A. Assay of catalases and peroxidases. Met. Enz. 1955; 2: 764–775. http://dx.doi.org/10.1016/S0076-6879(55)02300-8

43. Machado S, Paulsen GM. Combined effects of drought and high temperature on water relations of wheat and sorghum. Plant Soil. 2001; 233: 179–187. https://doi.org/10.1023/A:1010346601643

44. Kumar N, Poddar A, Shankar V, Ojha CSP, Adeloye AJ. Crop water stress index for scheduling irrigation of Indian mustard (*Brassica juncea*) based on water use efficiency considerations. J Agr Crop Sci. 2020; 206: 148–159. https://doi.org/10.1111/jac.12371

45. Caballero R, Goicoechea E, Hernaiz P. Forage yields and quality of common vetch and oat sown at varying seeding ratios and seeding rates of vetch. Field Crops Res. 1995; 41: 135–140. https://doi.org/10.1016/0378-4290(94)00114-R

46. Mafakheri A, Siosemardeh A, Bahramnejad B, Struik P, Sohrabi Y. Effect of drought stress and subsequent recovery on protein, carbohydrate contents, catalase and peroxidase activities in three chickpea (*Cicer arietinum*) cultivars. Aus J Crop Sci. 2011; 5: 1255–1260.

47. Pampana, S., Arduini, I., Andreuccetti, V., Mariotti, M. Fine-tuning N fertilization for forage and grain production of barley–field bean intercropping in Mediterranean environments. Agr 2022; 12: 1–20. https://doi.org/10.3390/agronomy12020418

48. Weisany W, Raei Y, Pertot I. Changes in the essential oil yield and composition of dill (*Anethum graveolens* L.) as response to arbuscular mycorrhiza colonization and cropping system. Ind Crops Pro. 2015; 77: 295–306. http://dx.doi.org/10.1016%2Fj.indcrop.2015.09.003

49. Machiani MA, Javanmard A, Morshedloo MR, Maggi F. Evaluation of competition, essential oil quality and quantity of peppermint intercropped with soybean. Ind Cro Pro. 2018; 111: 743–754. https://doi.org/10.1016/J.INDCROP.2017.11.052

50. Maffei M, Mucciarelli M. Essential oil yield in peppermint/soybean strip intercropping. Field Crops Res. 2003; 84: 229–240. http://dx.doi.org/10.1016/S0378-4290(03)00092-3

51. Almeselmani M, Deshmukh P, Sairam RK, Kushwaha S, Singh T. Protective role of antioxidant enzymes under high temperature stress. Plant Sci. 2006; 171: 382–388. https://doi.org/10.1016/j.plantsci.2006.04.009

52. Luhova L, Lebeda A, Hedererová D, Pec P. Activities of amine oxidase, peroxidase and catalase in seedlings of Pisum sativum L. under different light conditions. Plant Soil Env. 2003; 49: 151–157. http://dx.doi.org/10.17221/4106-PSE

53. Masoumi H, Darvish F, Daneshian J, Normohammadi G, Habibi D. Effects of water deficit stress on seed yield and antioxidants content in soybean (*Glycine max* L.) cultivars. Afr J Agr Res. 2011; 6: 1209–1218. https://doi.org/10.5897/AJAR10.821

54. Nair AS, Abraham T, Jaya D. Studies on the changes in lipid peroxidation and antioxidants in drought stress induced cowpea (*Vigna unguiculata* L.) varieties. J Env Bio. 2008; 29: 689–691. PMID: 19295066

55. Yadav R, Bhushan C. Effect of moisture stress on growth and yield in rice genotypes. Indian J Agr Res. 2001; 35: 104–107.

56. Nxele X, Klein A, Ndimba B. Drought and salinity stress alters ROS accumulation, water retention, and osmolyte content in sorghum plants. South Afr J Bot. 2017; 108: 261–266. https://doi.org/10.1016/j.sajb.2016.11.003

57. Agegnehu G. Ghizaw A. Sinebo W. Yield performance and land-use efficiency of barley and faba bean mixed cropping in Ethiopian highlands. Eur J Agr. 2006; 25: 202–207. http://dx.doi.org/10.1016/j.eja.2006.05.002

58. De la Fuente EB, Suárez SA, Lenardis AE, Poggio SL. Intercropping sunflower and soybean in intensive farming systems: evaluating yield advantage and effect on weed and insect assemblages. NJAS-Wag J Life Sci. 2014; 70: 47–52. https://doi.org/10.1017/wsc.2021.17

59. Amossé C, Jeuffroy MH, Mary B, David C. Contribution of relay intercropping with legume cover crops on nitrogen dynamics in organic grain systems. Nut Cyc Agr. 2014; 98: 1–14. https://dx.doi.org/10.1007/s10705-013-9591-8

60. Känkänen H, Eriksson C. Effects of undersown crops on soil mineral N and grain yield of spring barley. Eur J Agr. 2007; 27: 25–34. http://dx.doi.org/10.1016%2Fj.eja.2007.01.010

61. Luhmer K, Blum H, Kraska T, Döring T, Pude R. Poppy (*Papaver somniferum* L.) intercropping with Spring Barley and with White Clover: benefits and competitive effects. Agr. 2021; 11: 948. https://doi.org/10.3390/agronomy11050948

62. Latati M, Dokukin P, Aouiche A, Rebouh NY, Takouachet R, Hafnaoui E, Hamdani FZ, Bacha F, Ounane SM. Species interactions improve above-ground biomass and land use efficiency in intercropped wheat and chickpea under low soil inputs. Agr. 2019; 9: 765. https://doi.org/10.3390/agronomy9110765

63. Iliadis C. Evaluation of six chickpea varieties for seed yield under autumn and spring sowing. J Agr Sci. 2001; 137: 439–444.

64. Malhi SS. Improving crop yield, N uptake and economic returns by intercropping barley or canola with pea. Agr Sci. 2012; 8: 1023–1033. http://dx.doi.org/10.4236/as.2012.38124

65. Sadeghpour A, Jahanzad E, Lithourgidis A, Hashemi M, Esmaeili A, Hosseini M. Forage yield and quality of barley-annual medic intercrops in semi-arid environments. Int J Plant Pro. 2014; 8: 77–89.

66. Hauggaard-Nielsen H, Ambus P, Jensen ES. Interspecific competition, N use and interference with weeds in pea–barley intercropping. Field Crops Res. 2001; 70: 101–109. https://doi.org/10.1016/S0378-4290(01)00126-5

67. Hamzei J, Seyedi M. Evaluation of barley (*Hordeum vulgare*) and chickpea (*Cicer arietinum*) intercropping systems using advantageous indices of intercropping under weed interference conditions. Dan. Zeraat. 2014; 5, 1–12.

68. Fan Z, Chai Q, Yu A, Zhao C, Yin W, Hu F, Chen, G, Cao W. Coulter JA. Water and radiation use in maize–pea intercropping is enhanced with increased plant density. Agr J. 2020; 112: 257–273. https://doi.org/10.1002/agj2.20009

69. Andrade JF, Cerrudo A, Rizzalli RH. Monzon JP. Sunflower–soybean intercrop productivity under different water conditions and sowing managements. Agr J. 2012; 104: 1049–1055. https://doi.org/10.2134/agronj2012.0051

70. Megawer EA, Sharaan A, El-Sherif A. Effect of intercropping patterns on yield and its components of barley, lupine or chickpea grown in newly reclaimed soil. Egy J App Sci. 2010; 25: 437–452.

